# Glucocorticoids Unmask Silent Non-Coding Genetic Risk Variants for Common Diseases

**DOI:** 10.1101/2021.12.01.470787

**Authors:** Thanh Thanh Le Nguyen, Huanyao Gao, Duan Liu, Zhenqing Ye, Jeong-Heon Lee, Geng-xian Shi, Kaleigh Copenhaver, Lingxin Zhang, Lixuan Wei, Jia Yu, Cheng Zhang, Hu Li, Liewei Wang, Tamas Ordog, Richard M. Weinshilboum

## Abstract

Understanding the function of non-coding genetic variants represents a formidable challenge for biomedicine. We previously identified genetic variants that influence gene expression only after exposure to a hormone or drug. Using glucocorticoid signaling as a model system, we have now demonstrated, in a genome-wide manner, that exposure to glucocorticoids triggered disease risk variants with previously unclear function to influence the expression of genes involved in autoimmunity, metabolic and mood disorders, osteoporosis and cancer. Integrating a series of pharmacogenomic and pharmacoepigenomic datasets, we identified the *cis*-regulatory elements and 3-dimensional interactions underlying the ligand-dependent associations between those genetic variants and distant risk genes. These observations increase our understanding of mechanisms of non-coding genetic variant-chemical environment interactions and advance the fine-mapping of disease risk and pharmacogenomic loci.

## Main

A large number of genetic sequence variants associated with human disease have been discovered, but the task of understanding the function of those variants remains challenging since most of them map to non-coding regions of the genome^1, 2^. One approach to address this challenge has been to associate these variants with gene transcription, identifying so-called expression quantitative trait loci (eQTLs). Large-scale studies such as the Genotype-Tissue Expression (GTEx) Project has significantly improved our understanding of steady-state eQTLs across different tissues^3^. However, it is increasingly understood that eQTLs can be context-dependent or dynamic^4, 5^. For example, eQTL function can be modulated by pathogens^6^ or cellular differentiation^7^. Recently, we^8–10^ and others^11, 12^ have observed a series of uniquely “pharmacologic” eQTLs, hereafter referred to as “pharmacogenomic (PGx)-eQTLs”, for which eQTL behavior is elicited or significantly amplified in the presence of a drug or hormone. These dynamic eQTLs not only explain novel functions of non-coding variants but also provide valuable insight into molecular mechanisms underlying gene × environment interactions, interactions which could play important roles in complex disease pathophysiology^13^.

This study aimed to interrogate mechanistically, in a genome-wide manner, the interaction between non-coding genetic variants and the pharmacological environment for disease risk and variation in drug response by characterizing PGx-eQTLs with a series of “pharmaco-omic” datasets. Using an important pharmacological target, the glucocorticoid receptor (GR), as a study model, we were able to uncover a series of genome-wide PGx-eQTLs and obtain novel insight into how glucocorticoids might interact with genetic risk variants to predispose individuals to a wide range of pathology involving glucocorticoid signaling. Specifically, our study design (**Fig. 1a**) began with the identification of PGx-eQTLs using genome-wide single-nucleotide polymorphisms (SNPs) and RNA-seq and GR-targeted ChIP-seq before and after exposure to glucocorticoids in 30 immortalized human lymphoblastoid cell lines (LCLs) of differing genomic backgrounds. To validate drug-dependent effects, we treated these cells with cortisol, an endogenous GR agonist, and the drug CORT108297 (C297), a selective GR modulator which, in our studies, displayed antagonist properties when administered with cortisol and partial agonist activity by itself. Cortisol and its sister compounds are used routinely in the clinic to treat immunity-related diseases^14–16^, while C297 is a selective GR modulator currently being tested for the treatment of post-traumatic stress disorder (trial NCT04452500) and Alzheimer’s disease (trial NCT04601038). We then applied a series of epigenomic techniques including integrative chromatin state prediction (ChromHMM), a massively parallel reporter gene assay (STARR-seq) +/- drugs, and 3D chromatin conformation capture targeting *cis*-regulatory elements bound by the active enhancer- and promoter-associated histone mark acetylated histone H3 lysine 27 (H3K27ac HiChIP) +/- drugs. The integration of these datasets made it possible for us to determine underlying mechanism(s) and to generate additional evidence for associations underlying the observed PGx-eQTLs. Finally, we could then identify which of the discovered PGx-eQTLs might help to explain disease risk mechanisms by overlapping our SNPs with significant signals from publicly available genome-wide and phenome-wide association studies (GWAS and PheWAS).

**Fig. 1.**
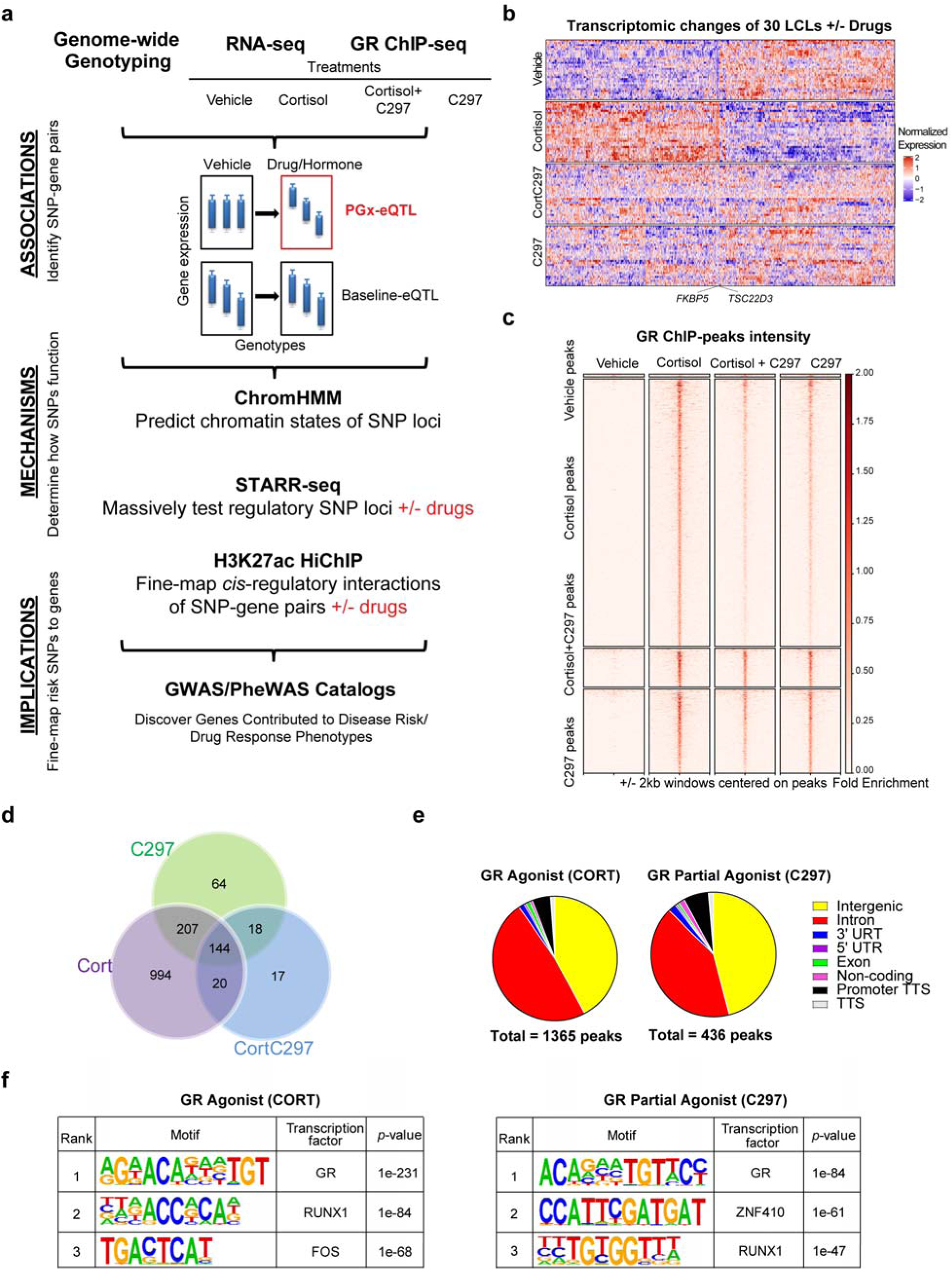
Conceptual Framework for the Study. (**a**) Experimental design used in this study. ChromHMM is a software used to annotate chromatin states from epigenomic data. STARR-seq stands for self-transcribing active regulatory region sequencing. Cortisol is an agonist, C297 is a modulator which acts as an antagonist when administered together with cortisol and a partial agonist by itself. (**b**) Heatmap of differentially expressed genes (FDR < 0.05) after four drug treatments across 30 LCLs showing drug-dependent patterns of gene regulation. Rows represent individual cell lines, and columns represent individual genes. (**c**) GR-targeted ChIP peak intensity after normalization to input across the 4 drug conditions used shows drug-dependent patterns similar to those of RNA-seq. (**d**) Overlap of GR-targeted ChIP-peaks across 4 drug conditions shows that C297 also acts as a partial agonist since the majority of C297-induced GR peaks overlapped with cortisol-induced peaks. (**e**) The distribution of GR-targeted ChIP peaks shows major enrichment in intronic and intergenic regions. (**f**) The top three de novo motifs identified by HOMER for GR-targeted ChIP-seq peaks after cortisol or C297 treatment demonstrates peak specificity to GR.

## Results

### Genome-wide discovery of glucocorticoid-modulated PGx-eQTLs in human LCLs

A total of 120 transcriptomic profiles were obtained under 4 drug conditions -- cortisol, C297, both together and vehicle control for 30 LCLs (**Supplementary Table 1**), as well as genome-wide GR-binding profiles under the same drug exposure conditions for a randomly selected LCL. The global changes of these profiles were characterized to ensure that the responses were glucocorticoid-specific. Because LCLs do not express or express only a very low level of mineralocorticoid receptor^4^, another endogenous receptor to which cortisol can bind, the treatment effects that we observed were GR-specific. We found that cortisol regulated the expression of 1361 genes across these 30 cells (FDR < 0.05) including GR canonical genes such as *FKBP5* and *TSC22D3*, whereas only 26 genes remained differentially expressed when C297 was added in combination with cortisol. Furthermore, C297 also acted as a partial agonist since, when tested alone, it upregulated a group of GR-target genes but to a lesser extent than cortisol, genes that included *FKBP5* and *TSC22D3* (see **Fig. 1b** for a visualization of drug-dependent transcriptomic patterns and **Supplementary Data 1** for a complete list of differentially expressed genes).

Similar drug-dependent patterns were observed in the GR-targeted ChIP-seq assays. Specifically, cortisol induced 1365 peaks and C297 induced 436 peaks at a 0.01 FDR threshold. These values were reduced to 200 peaks when the two drugs antagonized each other (**Figure 1c-d**). Approximately 85% of the C297-peaks overlapped with cortisol-peaks, suggesting that C297 is a partial agonist, as shown in the RNA-seq data (**Figure 1d**). In terms of distribution, GR peaks induced by either drug mapped predominately within intergenic and intronic regions, consistent with previous knowledge of GR function^17^ (**Figure 1e**). The total number of peaks was comparable to values obtained previously during GR ChIP-seq for LCLs using different GR antibodies and ligands^18^. We also demonstrated that these peaks were of high specificity, since *de novo* motif analysis showed that the peaks for both cortisol and C297 were most highly enriched in GR binding motifs (*P* = 10^-231^ for Cortisol, *P* = 10^-84^ for C297) (**Figure 1f**). After demonstrating that glucocorticoids displayed robust effects on RNA-seq and ChIP-seq, we set out to identify drug exposure-dependent eQTLs.

To leverage statistical power for our eQTL analysis, we narrowed the number of SNP-gene pairs by selecting only SNPs and genes that passed stringent quality control criteria. Specifically, 1.3 million genotyped SNPs were filtered based on Hardy-Weinberg Equilibrium (*P* > 0.001), genotyping call rates of more than 95%, and a minor allelic frequency of 0.18 to retain the probability of at least 1 cell line with the minor allelic genotype, resulting in 808,875 SNPs. We then identified SNPs within *cis* distances of +/- 200kb from genes, resulting in 433,272 SNPs. RNA-seq raw reads were normalized using conditional quantile normalization. All genes that mapped to sex chromosomes were excluded. Only genes with raw counts of ≥32 in at least one treatment and half of the LCLs were considered for eQTL analysis. All treatment conditions were then normalized to vehicle by genotypes to remove genotype effects at baseline and to adjust for cell line-dependent factors such as sex and age. We then selected SNPs that mapped within or near (+/- 500bp) GR binding sites by overlapping SNPs with the GR-targeted ChIP-seq data to focus on those most likely to interfere directly with GR binding and signaling, acknowledging that relevant SNPs with different possible mechanisms of action (e.g., SNPs within binding sites for downstream transcription factors regulated by GR) would be missed. As a result, we finally included a total of 1838 SNP-gene pairs for cortisol and 572 for C297 in the eQTL analysis.

Using the approach described in the preceding paragraph, we identified 102 cortisol-dependent and 32 C297-dependent *cis* PGx-eQTL SNP-gene pairs (*P* < 0.05) (**Fig. 2a-b, Extended Fig. 1**), the majority of which lost their cortisol-dependent eQTL status when exposed to C297 and cortisol together (*P* > 0.05), thus demonstrating the drug-dependent properties of this type of eQTL (**Fig 2e)**. Furthermore, these PGx-eQTLs were not significant baseline eQTLs based on data from 174 LCLs deposited in the GTEx database (*P* > 0.05) (**Fig. 2c-d**). We then sought to determine the molecular mechanisms for these SNPs. We observed that the SNPs themselves either mapped within known GR binding motifs, were in tight linkage disequilibrium with SNPs within GR motifs, or were distant from GR motifs (**Fig. 2f;** see **Supplementary Data 2** for details for each SNP). However, in all cases, they could still influence GR-dependent transcriptional activity, as later confirmed by massively parallel reporter gene assay.

**Fig. 2.**
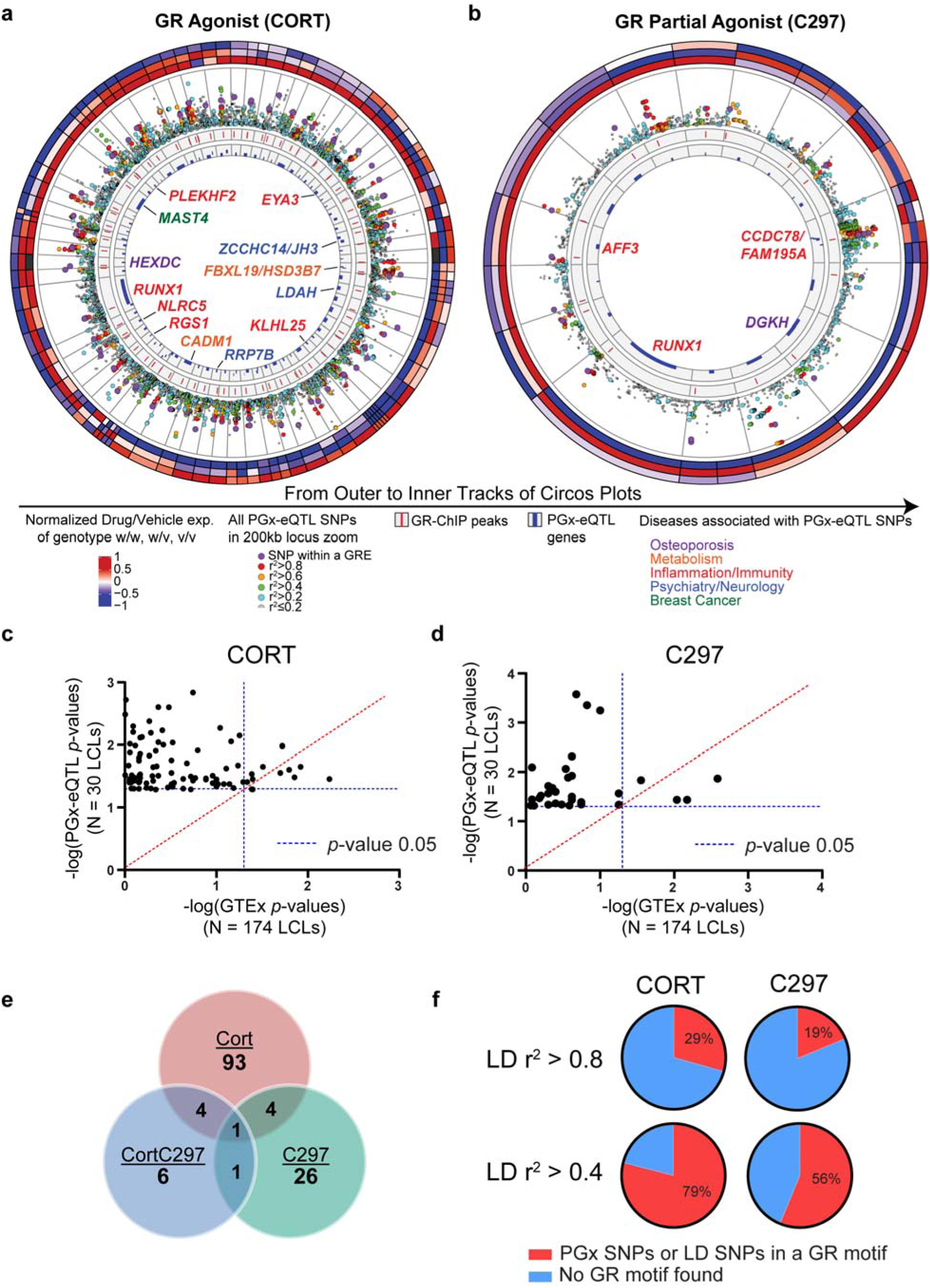
Discovery of GR-modulated PGx-eQTLs in human LCLs. (**a**-**b**) Circos plots depicting the GR-dependent PGx-eQTLs identified in 30 LCLs with results for cortisol (CORT) and the partial agonist (C297) shown in (**a**) and (**b**), respectively. The three outermost circles represent relative gene expression values (drug/vehicle) for each genotype, with each column depicting a PGx SNP-gene pair. The inner tracks are explained in the figure. The track with locus zoom plots m shows that SNPs within a GR binding site were generally more significantly associated with eQTL genes than other SNPs within the 200kb window of a gene. (**c**) & (**d**) *P*-Values for eQTL analyses from our study using 30 LCLs (Y axis) vs *P*-Values for eQTL analyses using 174 GTEx LCL samples. The majority of the PGx-eQTLs identified in the present study were not significant in GTEx even with larger sample sizes. (**e**) The number of SNP-gene pairs identified for each drug condition and their overlap across conditions. The majority of PGx-eQTL SNP-gene pairs after cortisol or C297 treatment no longer existed after antagonism (CortC297) was introduced. (**f**) Percentages of identified PGx-eQTL SNPs that mapped within a known GR binding motif or in tight LD with SNPs within a GR motif according to HaploReg V4.0 database.

### Glucocorticoid-modulated PGx-eQTLs most often mapped to enhancers with looping properties

Using ChromHMM, a software that models chromatin signatures with a multivariate Hidden Markov Model to annotate the putative regulatory function of the noncoding genome using epigenomic information^19^, we integrated 15 LCL epigenomic datasets from the Encyclopedia of DNA Elements (ENCODE) portal^20^ with our GR-targeted ChIP-seq data (see **Supplementary Data 3** for information on datasets). The 25-predicted chromatin states obtained were then annotated based on combinatorial and spatial patterns of chromatin marks^21^ (**Fig. 3a, Extended Fig.2, Supplementary Table 2**). These states could be categorized into four broad categories: (1) Promoter, (2) Enhancer, (3) Transcribed, and (4) Repressive/Repetitive/Unknown. Overlapping of GR-dependent PGx-eQTLs with these chromatin states demonstrated that PGx-eQTLs were enriched in a variety of states but predominantly in enhancers, with the primary site of enrichment being long-range enhancers that displayed promoter-looping properties (**Fig. 3b-c, Supplementary Table 3;** see **Supplementary Data 2** for details on each SNP). Specifically, 81% of cortisol-modulated PGx-eQTLs mapped to enhancer-related states and 41% mapped to enhancers with predicted looping properties. For C297-modulated PGx-eQTLs, 68% mapped to enhancer-related states and 65% mapped to enhancers with predicted looping properties.

**Fig. 3.**
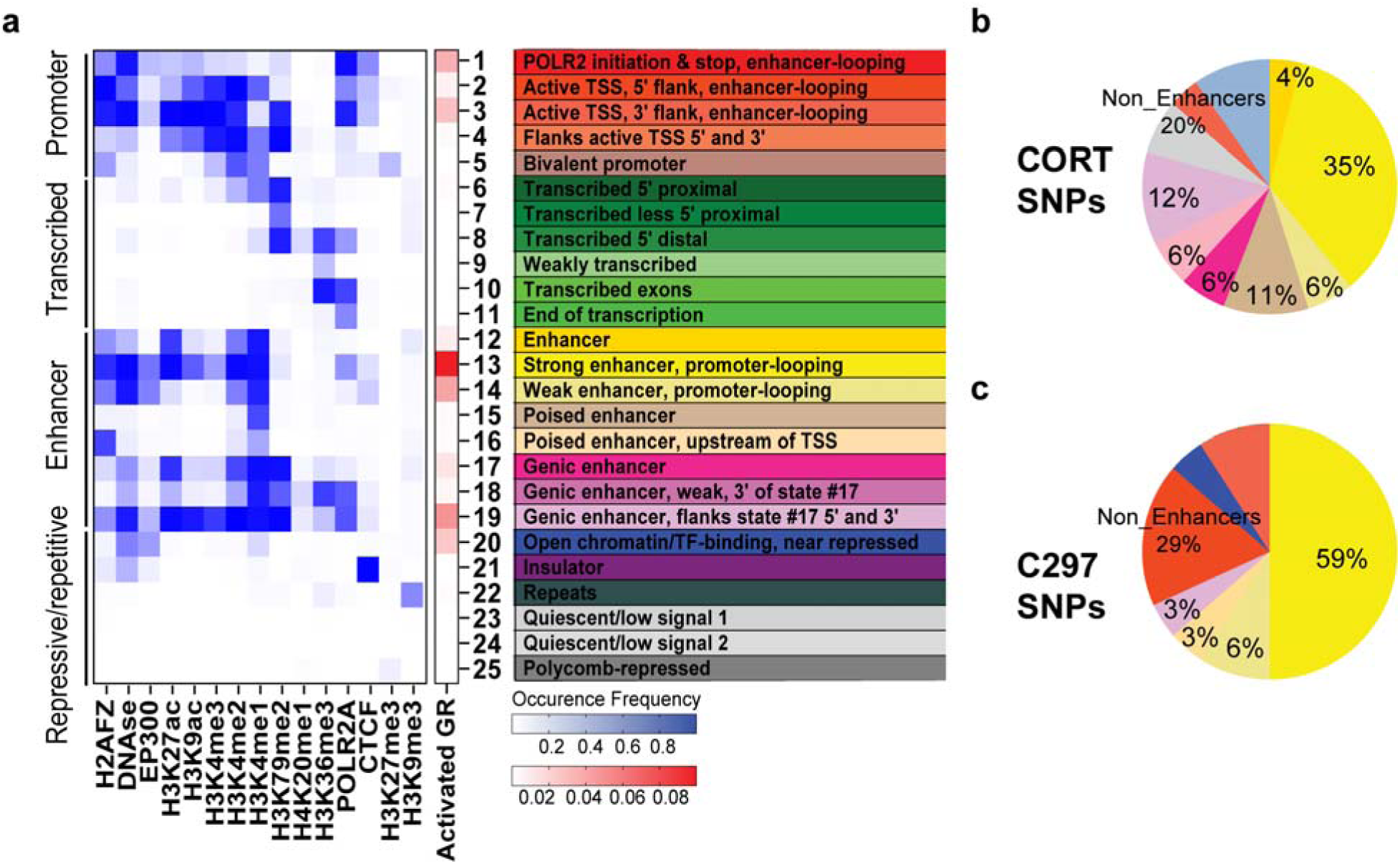
Prediction of chromatin states for GR-modulated PGx-eQTLs. (**a**) 25 LCL chromatin states predicted from the occupancy of 15 epigenetic marks for the reference LCL from ENCODE and the enrichment of GR peaks within each state. Columns represent epigenetic marks. Rows represent the co-occurrence probability of epigenetic marks within a state. (**b**-**c**) Distribution of GR-dependent PGx-eQTL SNPs among different chromatin states, which have been color coded as in (**a**).

### Allele-dependent and drug-dependent properties of enhancer PGx-eQTLs were replicated across cell lines

Because the majority of the PGx-eQTL SNPs that we had identified mapped to enhancer regions, we next performed STARR-seq, a massively parallel reporter gene assay that can capture enhancer activity in a high-throughput fashion^22^, to verify the effect of these SNPs and drugs on transcriptional activity associated with the identified PGx-eQTLs (see **Fig. 4a** for a diagram of the experimental workflow). We first cloned PCR-generated PGx-eQTL locus fragments from the pool of genomic DNA from our 30 LCLs into the human STARR-seq vector. Those fragments covered the GR binding peaks (+/- 500bp) that contained the identified PGx-eQTL SNPs. We included a glucocorticoid-induced GR peak in the library that mapped to a strong enhancer region near the *FKBP5* promoter as a positive control (**Extended Fig. 3a**). We then transfected the STARR-seq libraries into LCLs and A549 cells, a lung cancer cell line in which GR genomic regulation has been studied extensively^17^, exposed the cells to cortisol or C297, extracted mRNA and enriched the targeted sequences by RT-PCR. As expected, the transcribed products showed a size of around 1300bp (**Extended Fig. 3b**). After sequencing the transcribed loci, we mapped them to the human genome and achieved a mapping rate of more than 90% across all samples. We then called variants across loci, filtered out indels, multi-allelic variants, variants with low counts, and retained a total of 94 of the originally identified SNP-gene pairs for differential analysis. Replications for each sample were highly correlated (r^2^ ≥ 0.99) (**Extended Fig. 3c).**

**Fig. 4.**
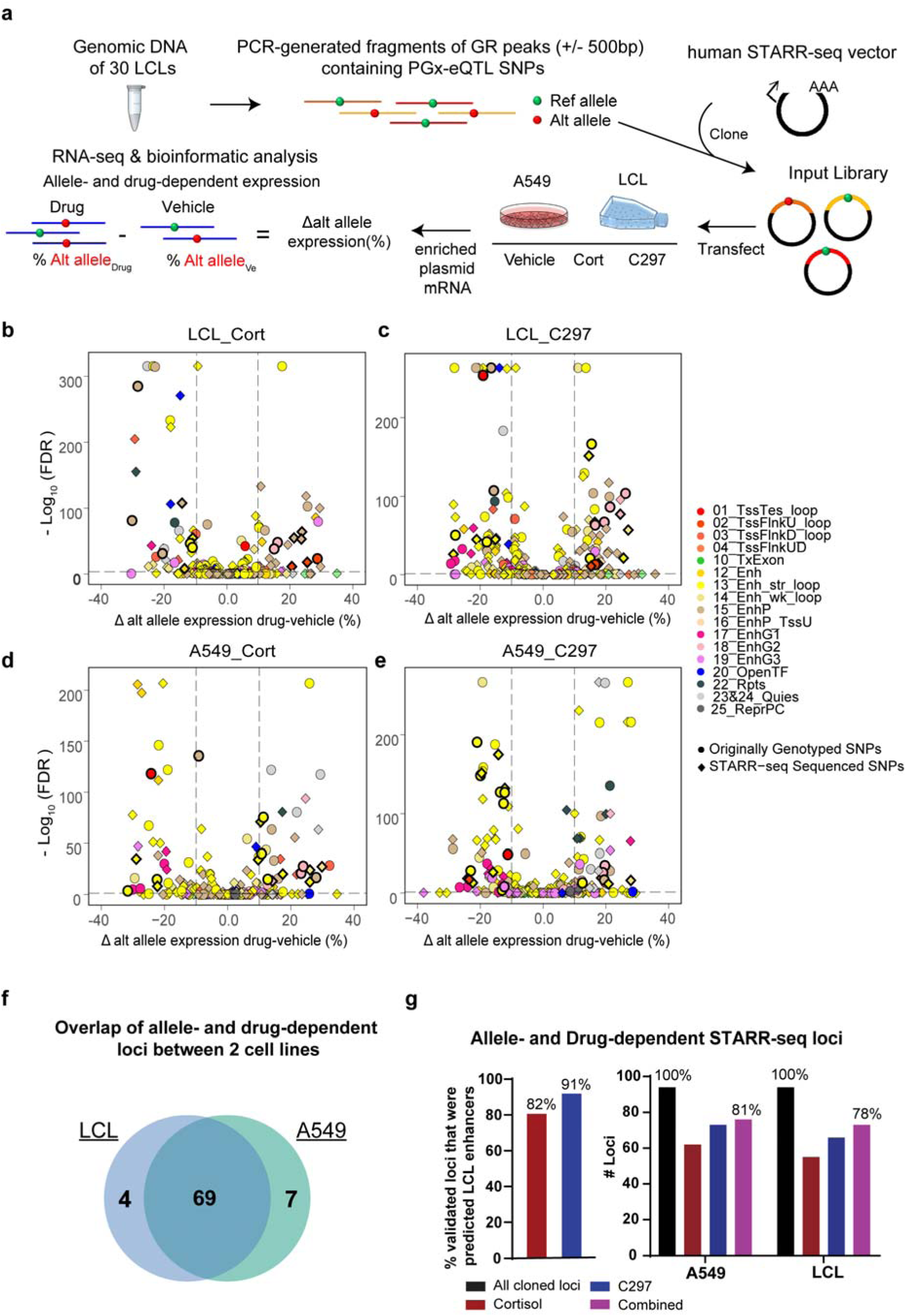
Testing allele- and drug-dependent effects of PGx-eQTLs with massively parallel reporter enhancer assay (STARR-seq). (**a**) Experimental workflow for STARR-seq: First, genomic DNA obtained from the 30 LCLs used in this study was collected. Second, PCR primers targeting GR-peaks +/-500bp that included identified PGx-eQTL SNPs were used to generate fragments containing loci with different genotypes. These fragments were then cloned into a human STARR-seq vector to generate input libraries. The libraries were then transfected into LCLs and A549 cells under different drug treatment conditions. mRNA was then extracted, enriched for transcripts transcribed from the STARR-seq vector, and prepared for next-generation sequencing and bioinformatic analyses to identify allele- and drug-dependent transcriptional activities. (**b-e**) Volcano plots showing PGx-eQTL loci where SNP-dependent and GR-dependent activities were detected. The Y axis represents –log 10 of FDR from Fischer’s Exact Test, the X axis represents change of alternative allele percentages after drug treatment. Circles represent originally genotyped SNPs, and squares represent SNPs sequenced in each STARR-seq locus. Each SNP was color coded by the chromatin state in which they resided. All loci that were later found to be associated with diseases that achieved statistical significance for SNP-dependent and GR-dependent transcriptional analysis are bolded in black. (**f**) High consistency between the two cell lines, LCL and A549, used in the STARR-seq assay in terms of loci that displayed allele- and drug-dependent properties. (**g**) STARR-seq results validated allele- and drug-dependent enhancer activities of identified PGx-eQTL loci.

Because all samples shared the same input library, we focused on analyzing differences of STARR-seq transcriptional activities among drug treatment conditions and alleles. Principal component analysis showed global differences among drug treatments, demonstrating drug effects on STARR-seq transcription (**Extended Fig. 3d**). To test for allele- and drug-dependent effects, we analyzed percentage differences of alternative alleles before and after drug treatments. In LCLs, the expression of 44 originally genotyped SNPs was allele-dependent after cortisol treatment, as were 55 after C297 treatment (FDR < 0.05, percentage change of alternative allele > 10%) (**Fig. 4b**). *De novo* SNPs identified by sequencing in STARR-seq demonstrated that 55 (cortisol) and 67 (C297) loci had allele- and drug-dependent transcriptional activity (**Fig. 4c)**. In A549 cells, expression of 52 originally genotyped SNPs was allele-dependent after cortisol treatment, and 61 after C297 treatment **(Fig. 4d)**. *De novo* SNPs identified by sequencing in STARR-seq demonstrated that 62 (cortisol) and 73 (C297) A549 cell loci had allele- and drug-dependent transcriptional activity (**Fig. 4e)**. These loci displayed high consistency between the two cell lines in which STARR-seq was applied (**Fig. 4f**). As anticipated, the majority of allele- and drug-dependent loci identified by STARR-seq were ChromHMM-predicted enhancers, validating up to 81% of the cloned PGx loci that we had identified (**Fig. 4g,** see **Supplementary Data 2** for details on each locus).

### PGx-eQTL SNP-gene pairs are connected by drug-dependent enhancer-enhancer and enhancer-promoter loops

To determine the nature of physical interactions between PGx SNP loci and eQTL genes, we applied H3K27ac HiChIP, an assay that can capture chromatin conformation of enhancer-promoter and enhancer-enhancer interactions in a high-resolution manner^23^, before and after cortisol exposure. First, to demonstrate that drug treatment in the HiChIP experiment was successful, qRT-PCR was conducted for *FKBP5*, a prototypical GR-targeted gene, using total RNA from the same cells before fixation. *FKBP5* was induced ∼6 fold after drug treatment, confirming drug treatment effect (**Fig. 5a**). After library preparation, H3K27ac ChIP efficiency was achieved at 0.21% to 0.42% of total input for vehicle and cortisol, respectively. Shallow sequencing confirmed that the percentage of PCR duplication was less than 0.01%, and that more than 40% of the fraction of reads represented interactions within the same chromosome (long-range *cis* interactions). Deep sequencing yielded 829,599,511 raw PE reads for vehicle (of which 88.1% were mapped), and 539,417,612 for cortisol (of which 87.4% were mapped). Using known ENCODE H3K27ac ChIP-peaks for LCLs for loop calling at an FDR threshold of 0.01, the number of called loops was 193107 for vehicle and 131926 for cortisol, which displayed high enrichment around ChIP-peaks (**Extended Fig. 4a**). These numbers are comparable with those reported for a publicly available H3K27ac HiChIP library for LCL GM12878 (185167 loops)^23^. With H3K27ac ChIP-peaks called directly from our data, the number of loops called for each sample was slightly higher but with less enrichment around the peaks. Therefore, we called loops using the previously generated ChIP-seq data to avoid false positive signals, understanding that we might miss some H3K27ac signals that were induced/repleted by cortisol, if they might exist. The percentage of intra-chromosomal interactions that spanned greater than 15kb in linear genomic distance, a measure of how well the library captured chromatin interactions between genomic loci, was 35.7% for Vehicle and 36.9% for cortisol, surpassing the manufacturer’s minimum benchmark of 25%. Percentages of valid interaction pairs located within known ChIP-seq peaks was 37.5% for Vehicle and 33.5% for cortisol, surpassing the manufacturer’s minimum benchmark of 15%.

**Fig. 5.**
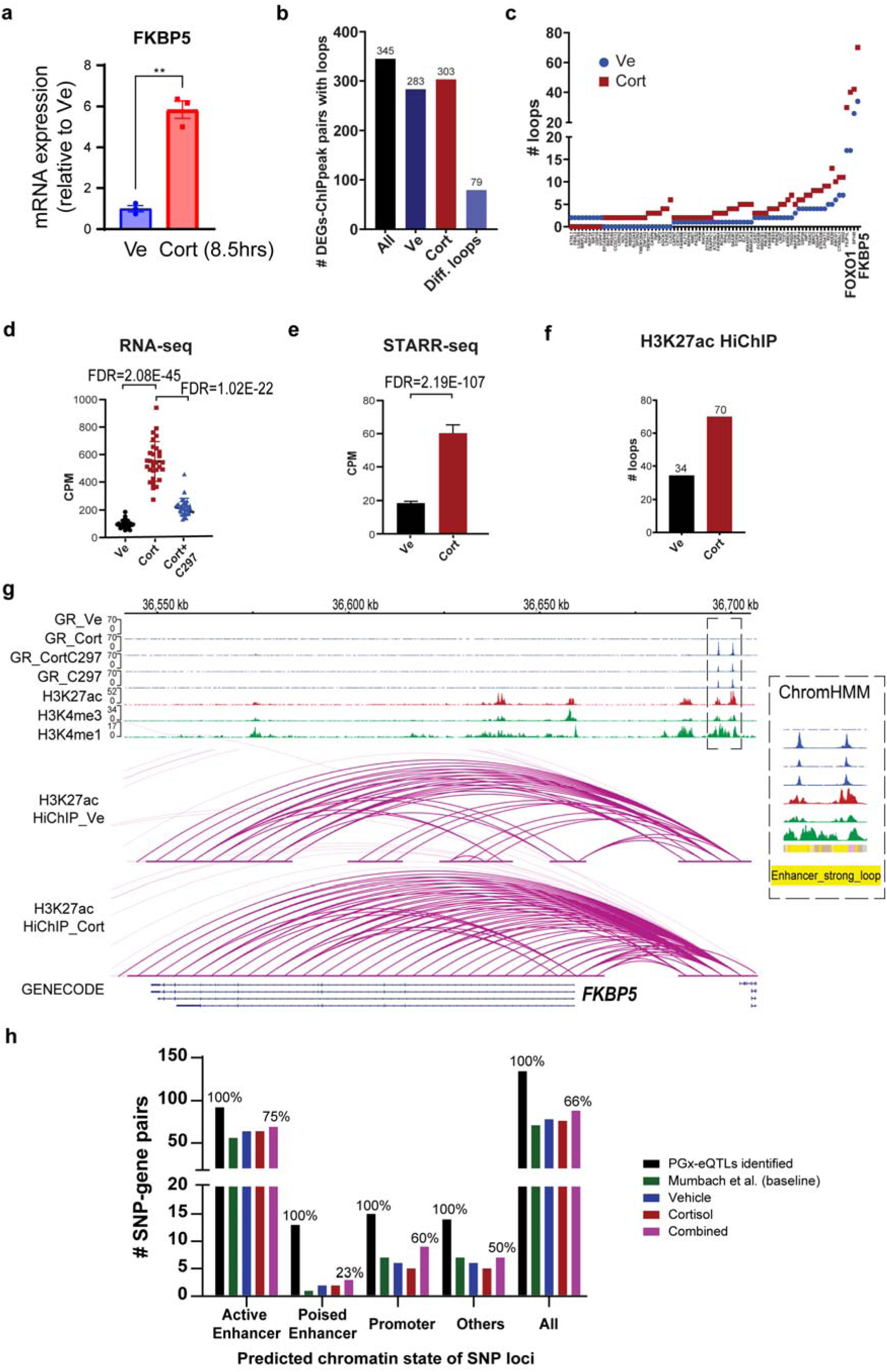
Generation and integration of H3K27ac HiChIP before and after glucocorticoid treatment with other datasets. **(a)** Validation of drug treatment effect for HiChIP samples as measured by qRT-PCR for *FKBP5*, a prototypical GR-targeted gene, using RNA extracted from the same cells. Statistical significance was evaluated with student t-test, achieving *P* < 0.005. Each dot represents a replicate. **(b)** Number of cortisol-regulated genes that were connected to a cortisol-induced GR ChIP-peak by a H3K27ac loop in different drug conditions. (**c**) Cortisol-regulated genes that were connected to one or more cortisol-induced GR ChIP-peaks by differential cortisol-regulated H3K27ac loop(s) (fold change > 1.5 or a change from 0 that is more than 1). The X axis shows gene names. (**d**) mRNA expression of *FKBP5* before and after drug treatment as determined by RNA-seq. CPM represents counts per million. (**e**) Transcriptional activity driven by the enhancer region upstream of *FKBP5* as measured by STARR-seq. (**f**) Number of HiChIP H3K27ac loops that connected GR-binding sites to the *FKBP5* gene. (**g**) Integrative Genomics Viewer (IGV) plots of 2 different GR-dependent enhancers over a distance of 50kb, which together regulated *FKBP5*. These two enhancers were predicted to be strong enhancers with looping properties by ChromHMM. (**h**) Number of PGx-eQTLs that displayed physical interactions between SNP loci (categorized by enhancer/non-enhancer states) and eQTL genes as demonstrated by H3K27ac HiChIP.

To determine whether H3K27ac loops changed after cortisol treatment and to what extent they might be correlated with functional outcome (differential gene expression), we integrated the cortisol-regulated RNA-seq, GR ChIP-seq and HiChIP datasets. Out of 1361 differentially expressed genes (DEGs), 345 had HiChIP loops connecting them to one or more cortisol-induced ChIP-seq peaks, and 79 DEGs had loops that were altered after drug treatment (defined as a fold change in number of loops > 1.5 or a change from no loops to a number > 1) (**Fig. 5b-c**). To help readers visualize this integrative approach, which was applied later to fine-map PGx-eQTL SNP-gene pairs, we showcased two positive controls for all of the datasets in **Fig 5d-g** and **Extended Fig 4b-d**. Specifically, RNA-seq showed that *FKBP5* mRNA expression was induced by cortisol (FDR*=*2.08E-45), an induction that was reversed by C297 (FDR*=*1.02E-22). This observation for gene expression correlated with the GR-binding patterns under the same drug conditions at the enhancer regions that mapped 50kb upstream of *FKBP5* (**Fig. 5d-e**). This region was predicted to be a strong enhancer with looping properties (**Fig. 5g).** After being tested in STARR-seq, it showed a strong induction in enhancer activity after drug treatment (FDR=2.19E-107) (**Fig. 5e)**. The number of H3K27ac HiChIP loops also increased 2-fold after cortisol treatment, connecting the GR-induced enhancers to *FKBP5*, transcriptionally regulating the gene (**Fig. 5f**). Similar observations were made for *FOXO1*, for which GR-modulated enhancer signals were integrated from four different regions over hundreds of kilobases, with a 2-fold change of HiChIP loops after drug treatment (**Extended Fig. 4b-d)**.

As expected, most of the SNPs that were connected with PGx-eQTL genes with H3K27ac loops mapped within ChromHMM-predicted active enhancers (**Fig. 5h**). Publicly available data^23^ showed that 71 of the PGx-eQTLs that we identified displayed H3K27ac connecting “loops” at baseline. We observed a similar number of connected PGx-eQTLs before and after cortisol treatments in our data (78 and 76, respectively) (**Fig. 5h)**, with 68 being constitutive loops connecting the PGx SNP loci and eQTL genes but requiring glucocorticoids to “unmask” their functional transcriptional regulation. In fewer cases, cortisol was shown to induce loops for 5 SNP-gene pairs or to repress loops for 6, either bringing together SNP-gene loci that otherwise would not have been in proximity or separating them (**Supplementary Data 2**). These mechanisms are consistent with previous observations obtained by Hi-C in A549 cells +/- glucocorticoid treatment^24^.

### Glucocorticoid-modulated PGx-eQTLs unmasked potential function of SNP loci previously associated with common diseases involving glucocorticoid signaling

In an unbiased search of GWAS/PheWAS databases (https://www.ebi.ac.uk/gwas/, https://pheweb.sph.umich.edu/, https://r4.finngen.fi/about), we found that twenty-five percent of the glucocorticoid-modulated PGx-eQTL SNPs that we identified had previously been associated with clinical phenotypes but usually without a clear underlying mechanism and often lacking clarity with regard to the gene or genes involved. These associations spanned many different disease and drug response categories including adverse response to corticosteroids, inflammation and immunity, osteoporosis, neuropsychiatric disorders and cancer (**Table 1;** see **Supplementary Data 2** for integrative annotation of each locus). For example, a mechanistically unexplained variant that had been associated with breast cancer risk, rs1697139^25^, was a cortisol-dependent PGx-eQTL for the Microtubule Associated Serine/Threonine Kinase Family Member 4 (*MAST4*) gene. Specifically, *MAST4* expression was repressed by cortisol in subjects with the G/G but not the A/A genotype (Adjusted *P*=0.0061), and that repression was reversed after antagonist treatment (Adjusted *P*=0.3446) (**Fig. 6a**). This SNP, in a genotype-dependent fashion, modulated a GR-responsive intergenic enhancer (FDR=1.62E-41) that “looped” across 40,000 base pairs to *MAST4*, transcriptionally regulating that gene (**Fig. 6b-c)**. Of interest, in breast cancer cell lines, GR bound to the same rs1697139 locus and dramatically repressed *MAST4* expression in MDA-MB-231 cells, a triple-negative breast cancer cell line, as well as cell lines for other breast cancer subtypes (**Fig. 6d-f**, **Extended Fig. 5a-f**). Although the function of *MAST4* in breast cancer is unknown, it may be a novel risk gene since its expression was significantly repressed in breast tumor tissue when compared with normal breast tissue (*P*<0.0001) (**Fig**. **6g)**. Furthermore, *MAST4* expression also appeared to be a predictor of treatment response since decreased expression of *MAST4* was associated with decreased relapse-free survival (*P*=1.6E-10) (**Fig. 6h**). These observations agree with the direction of the SNP-phenotype association. Specifically, the rs1697139-G/G genotype was associated with decreased *MAST4* expression after exposure to cortisol, a hormone that promotes breast cancer heterogeneity and metastasis^26^, which might have led to increased risk for breast cancer observed in the original GWAS. Of interest is the fact that glucocorticoids were recently found to induce chemo-resistance in solid tumors by transcriptionally regulating another MAST family member, *MAST1*^27^, a protein which has the most homology with *MAST4* within this protein family^28^.

**Fig. 6.**
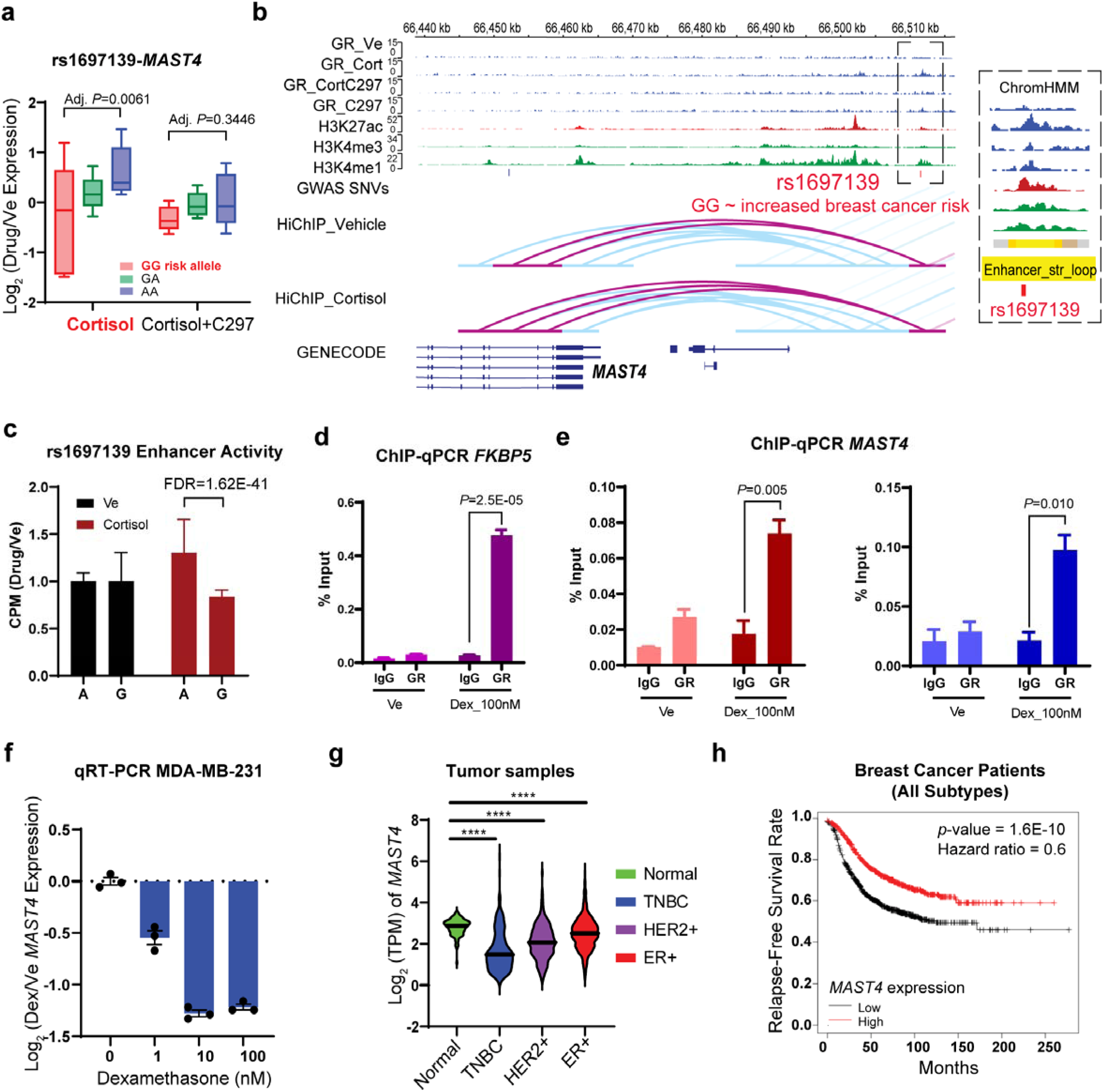
Example of GR-dependent PGx-eQTLs with functional implications for disease risk. **(a)** PGx-eQTL SNP-gene pairs for rs1697139-*MAST4*. Adjusted *P*-values from Tukey’s post-hoc multiple comparisons and represent differences between wildtype and variant genotypes. Cortisol induced the eQTL, and C297 antagonized the cortisol effect, normalizing eQTL expression across genotypes. (**b**) IGV plots of the PGx SNP-eQTL gene locus. Tracks for GR- targeted ChIP-seq in different drug conditions are colored in blue, which show similar drug-dependent pattern as expression data: Cortisol induced GR binding at SNP locus, and C297 antagonized the cortisol effect, reducing GR binding. H3K4me1 is a histone mark associated with enhancers. H3K4me3 is a histone mark associated with promoters. H3K27ac is a histone mark associated with active promoters and enhancers. For the H3K27ac HiChIP tracks, loops directly interacting with the PGx SNP locus are highlighted in pink and others in blue. (**c**) SNP- dependent and drug-dependent enhancer activity of the PGx locus as measured by STARR-seq. CPM stands for counts per million. (**d**) Positive control for ChIP-qPCR assays that tested GR binding at the *FKBP5* enhancer region in the TNBC cell line MDA-MB-231. (**e**) ChIP-qPCR assays that tested GR binding at the PGx locus for MAST4 in the TNBC cell line MDA-MB-231 using two different primers. *P*-values from student t-tests. (**f**) Dose-dependent repression of *MAST4* by glucocorticoids in the TNBC cell line MDA-MB-231. (**g**) Expression of *MAST4* in tumors from breast cancer patients in the TCGA database. **** *P*-values < 0.0001 by student t-tests. TPM stands for transcripts per million. (**h**) Kaplan-Meier curves for relapse-free survival rate in 1764 breast cancer patients predicted based on *MAST4* expression.

**Table 1.**
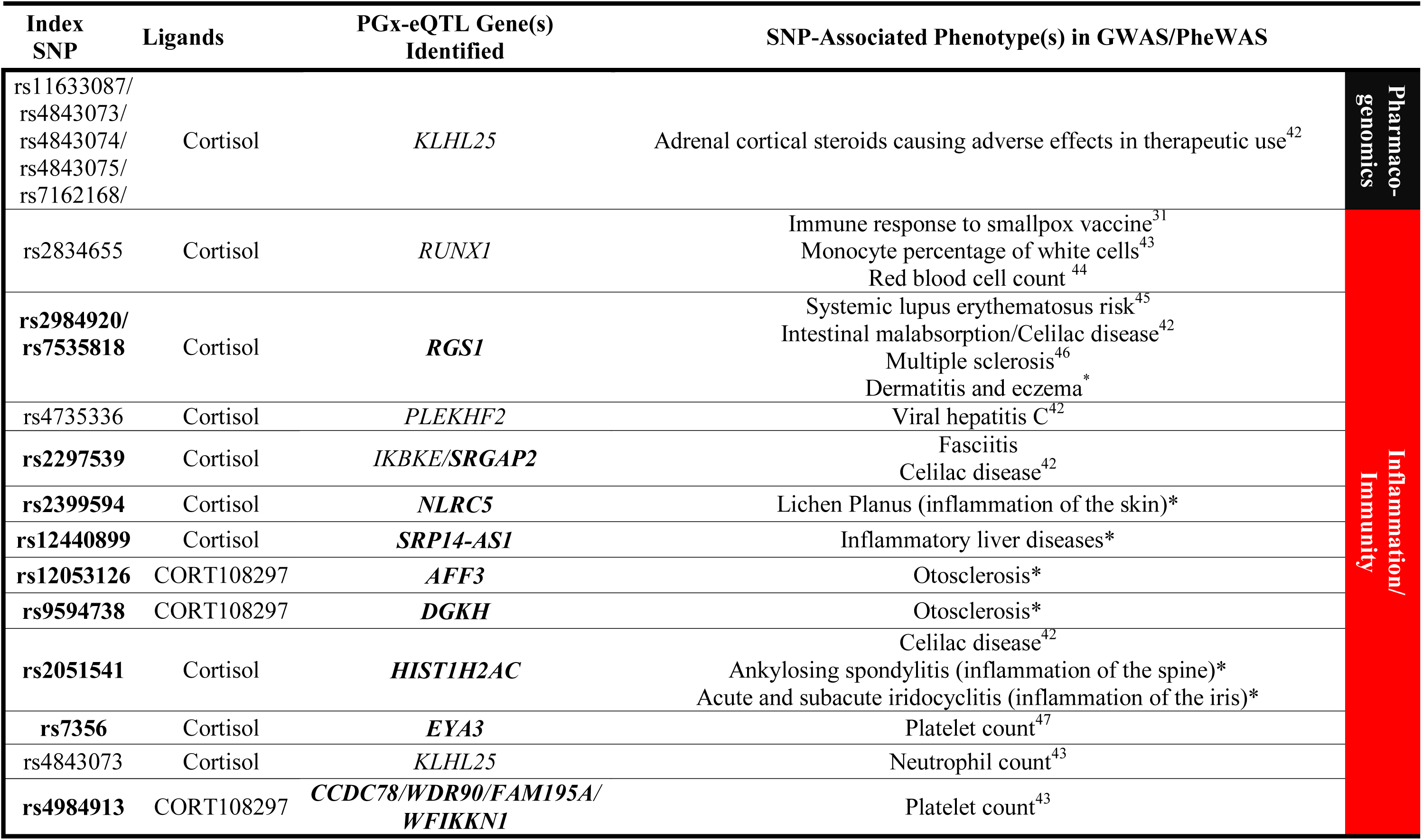
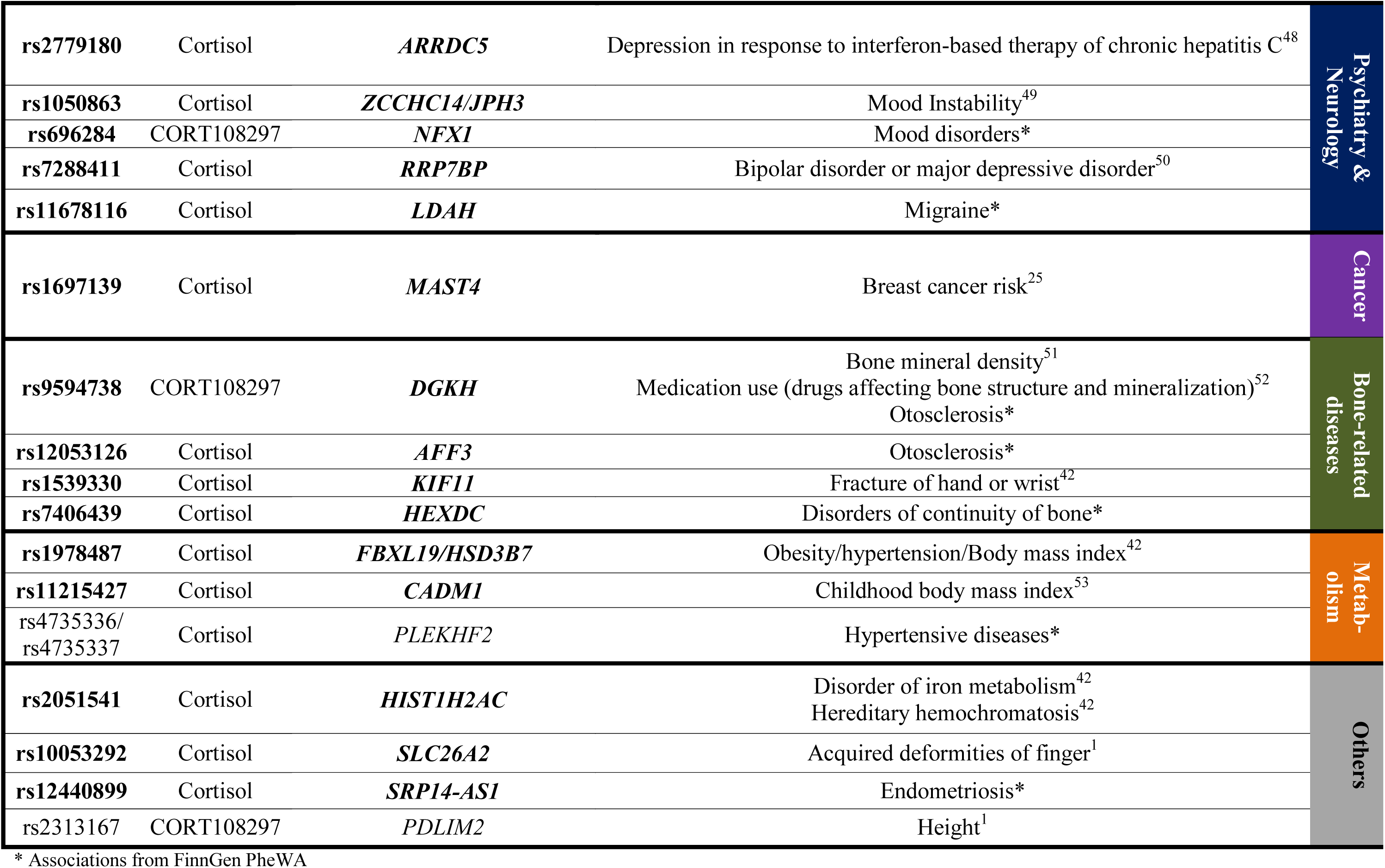
GR-modulated PGx-eQTLs which we identified that have been associated with a clinical phenotype by previous GWAS/PheWAS. SNP-gene pairs that were PGx-eQTLs (P < 0.05) but not eQTLs in GTEx (P > 0.05) are bolded

Beyond this illustrative example**, Table 1** lists a total of 30 disease risk loci that we found to behave as PGx-eQTLs. Of importance, glucocorticoids were either known risk factors or therapeutic agents used to treat most of these diseases, demonstrating a genetic risk × hormone/drug risk interaction in disease predisposition. For instance, glucocorticoids are potent immune-suppression and anti-inflammatory agents^14^. As a result, GR agonists are used clinically to treat a wide variety of immune-related diseases including those listed in **Table 1**–e.g., multiple sclerosis^15^ and systemic lupus erythematosus^16^. Corticosteroids are also known to increase platelet number and, as a result, are first-line therapy for immune thrombocytopenia^29^. In another example, we found that a SNP previously associated with response to vaccine and counts for different immune-related blood-cell types interacted with cortisol to influence the expression of RUNX Family Transcription Factor 1, *RUNX1* (**Extended Fig. 6a),** a major regulator of hematopoiesis^30^. Specifically, the *RUNX1* G/G genotype increased GR binding, correlating with an increase in GR-modulated enhancer activity, an intergenic enhancer that “looped” to *RUNX1*, resulting in decreased *RUNX1* expression (**Extended Fig. 6b-d**). The repression of *RUNX1* could result in a decrease of B cells^30^, which might help to explain, in part, why antibody levels after vaccination were decreased in individuals carrying the G/G genotype^31^. We also identified a SNP previously associated with lichen planus, an autoimmune condition that attacks cells of the skin and mucosus membranes, that interacted with cortisol to influence the expression of *NLRC5*, a key regulator of adaptive immune responses^32^.

In addition to inflammation and immunity, glucocorticoids also play an important role in osteoporosis^33^, mood disorders^34, 35^, cancer^26^ (as described above), and metabolism^36^, diseases that are also listed in **Table 1**. In an example involving the rs11678116 SNP and the Lipid-Droplet Associated Hydrolase (*LDAH*) gene, cortisol was shown to bring together SNP-gene loci that otherwise would not have been in proximity, “unmasking” the impact of rs11678116 on *LDAH* transcription (**Fig. 7a-b**). Specifically, the rs11678116 SNP created a GR binding motif (**Supplementary Data 2**), which led to decreased cortisol-responsive enhancer activity of the T/T genotype and decreased *LDAH* expression after cortisol treatment of subjects with the T/T but not the G/G genotype (**Fig. 7c**). Based on PheWAS results, rs11678116 was also associated with migraine (**Table 1**), a stress-sensitive condition for which cortisol is a biomarker^37^, providing an intriguing genotype-phenotype link for functional investigation given the elevated levels of cholesterol and triglycerides observed in migraine patients^38^. We also found that SNPs previously associated with body mass index and obesity interacted with cortisol to influence genes located as far as 150,000bp away such as *FBXL19*, an adipogenesis-controlling gene^39^, *HSD3B7*, a cholesterol metabolizing enzyme (**Extended Fig. 8a-b**), or *CADM1*, a gene that regulates body weight via neuronal modulation^40^. Taken together, these examples — all of which will require further study to functionally validate — demonstrate that glucocorticoid-dependent PGx-eQTLs identified in LCLs uncovered functional SNPs related not only to immune-related diseases but also diseases reflecting dysfunction of various other cell types. That is consistent with previous observations that eQTLs that are shared across tissues comprise a larger fraction of trait associations than do tissue-specific eQTLs^41^.

## Discussion

Despite the many associations with human disease that have been described for non-coding genetic variants, their functional interpretation remains a significant challenge^1, 2^. Furthermore, complex diseases are usually influenced by both genetic and environmental factors, which are difficult to interrogate mechanistically^13^. This manuscript addresses these two themes by mechanistic studies of a type of non-coding genetic variant with functions that are modulated by pharmacologic or physiologic chemical agents. Obviously, this study has limitations previously discussed that are inherent to the study of response-eQTLs^5^ – namely, limited cell types that are available as study models and limited power as a result of the resources required to generate datasets with and without chemical stimuli. We addressed these issues by studying combinations of treatment with agonists and antagonists to verify drug-dependent effects. We then provided additional layers of mechanistic evidence to support our observations but with the clear acknowledgement that we might fail to capture all relevant SNPs. We also validated selected examples with experiments in other relevant cell lines, coupled with the integration of available clinical data for functional interpretation.

By systematically fine-mapping genotype-phenotype interactions in which measurable environmental factors such as drug or hormone exposure were taken into account, we uncovered potential novel risk genes for a range of diseases in which the pharmacological or physiological stimuli played important roles. The function of these SNPs was usually “masked” in the absence of exposure to hormones or related drugs. As a result, exposure to these compounds could either initiate transcriptional activity between connected loci or elicit a conformational change in the epigenomic landscape to disrupt or bring the SNP locus into contact with distal gene(s) that were often relevant to the observed clinical phenotypes. Functional studies of genes discovered via this mechanism have already yielded novel insights into mechanisms of disease^8, 10^. As a result, this study has added a novel perspective to functional genomics by providing a mechanistic framework for additional studies of ligand-dependent “silent” non-coding genetic variants to advance the fine-mapping of disease risk and pharmacogenomic loci. Insights from those efforts could partly explain mechanisms underlying genetic and environmental susceptibility to common diseases or variation in response to drug therapy.

## Acknowledgments

This work was supported by the U.S. National Institute of General Medical Sciences (grant no. U19GM61388 to RWM and LWang and R01GM28157 to RMW), National Institute of Alcohol Abuse and Alcoholism (grant no. R01AA027486 to RWM), National Institute of Diabetes and Digestive and Kidney Diseases (grants no. R01DK126827 and R01DK058185 to TO), and the Mayo Research Foundation (to RWM). We thank the GTEx and ENCODE Consortiums for their generation of transcriptomic and epigenomic datasets used in this study. We also want to acknowledge the participants and investigators of the FinnGen study.

## Author contributions

Conceptualization: TTLN, DL, TO, RMW; Investigation: TTLN, DL, LZ; Data Curation: HG, KC, LWei, HL, CZ; Formal Analysis: HG, TTLN, ZY; Methodology: JHL, GS, TO; Resources: LWang, TO; Supervision: RMW, LWang, TO; Funding Acquisition: RMW and LWang; Visualization: TTLN, HG, KC; Data Interpretation: TTLN, DL, HG, JY, LWang, TO, RMW ; Writing – original draft: TTLN, RMW; Writing – review & editing: DL, HG, JHL, LZ, JY, TO, RMW

## Data and materials availability

All sequencing data was deposited on Gene Expression Omnibus under accession number GSE185941.

---

**Extended Fig.1.**
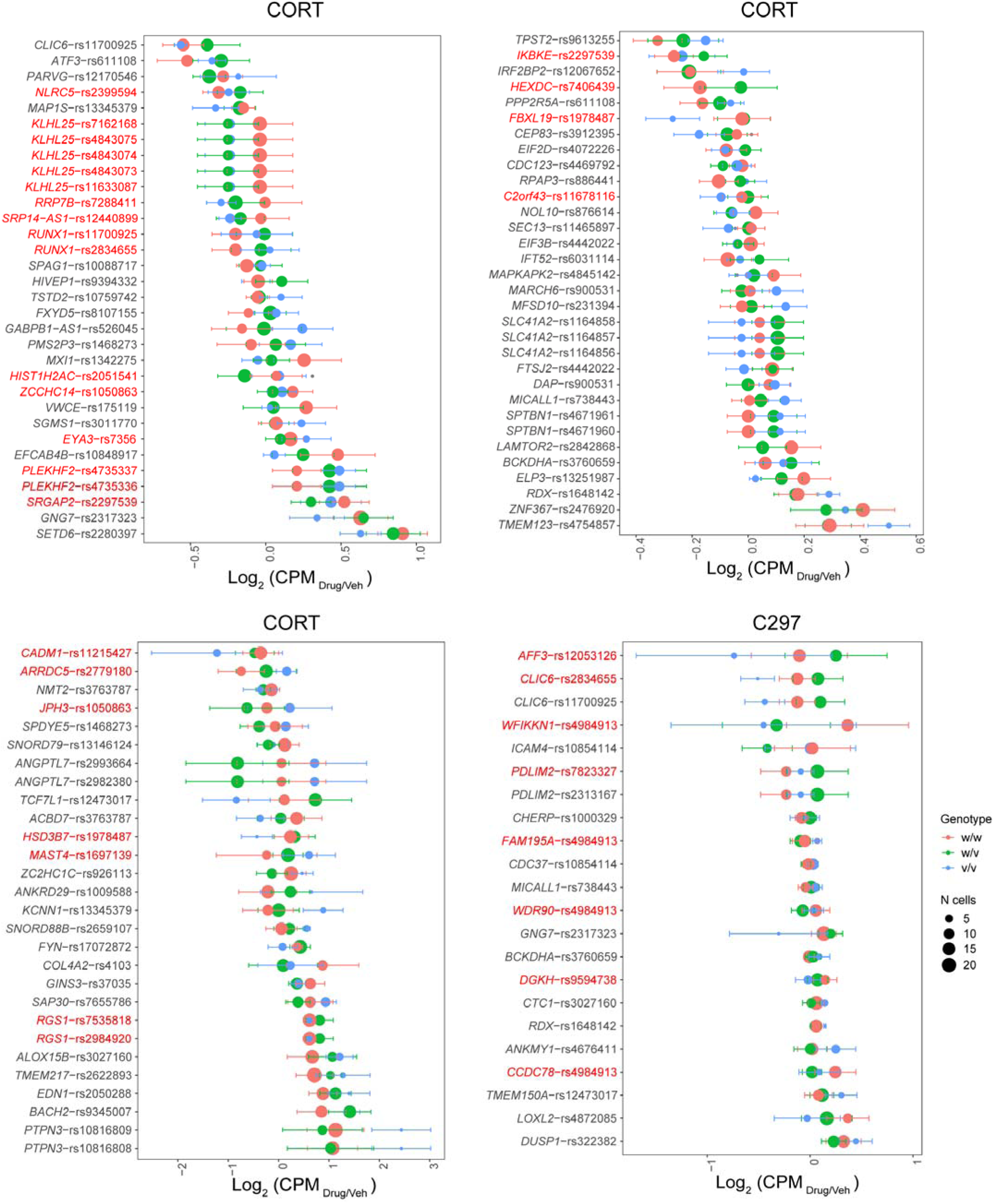
Identification of GR-dependent PGx-eQTLs. Dot plots depicting eQTL gene expression patterns for drug condition normalized to vehicle. Genotypes are color coded. Dot sizes represent the number of cell lines available for each genotype. PGx-eQTL SNPs that have previously been associated with a clinical phenotype are highlighted in red text.

**Extended Fig.2.**
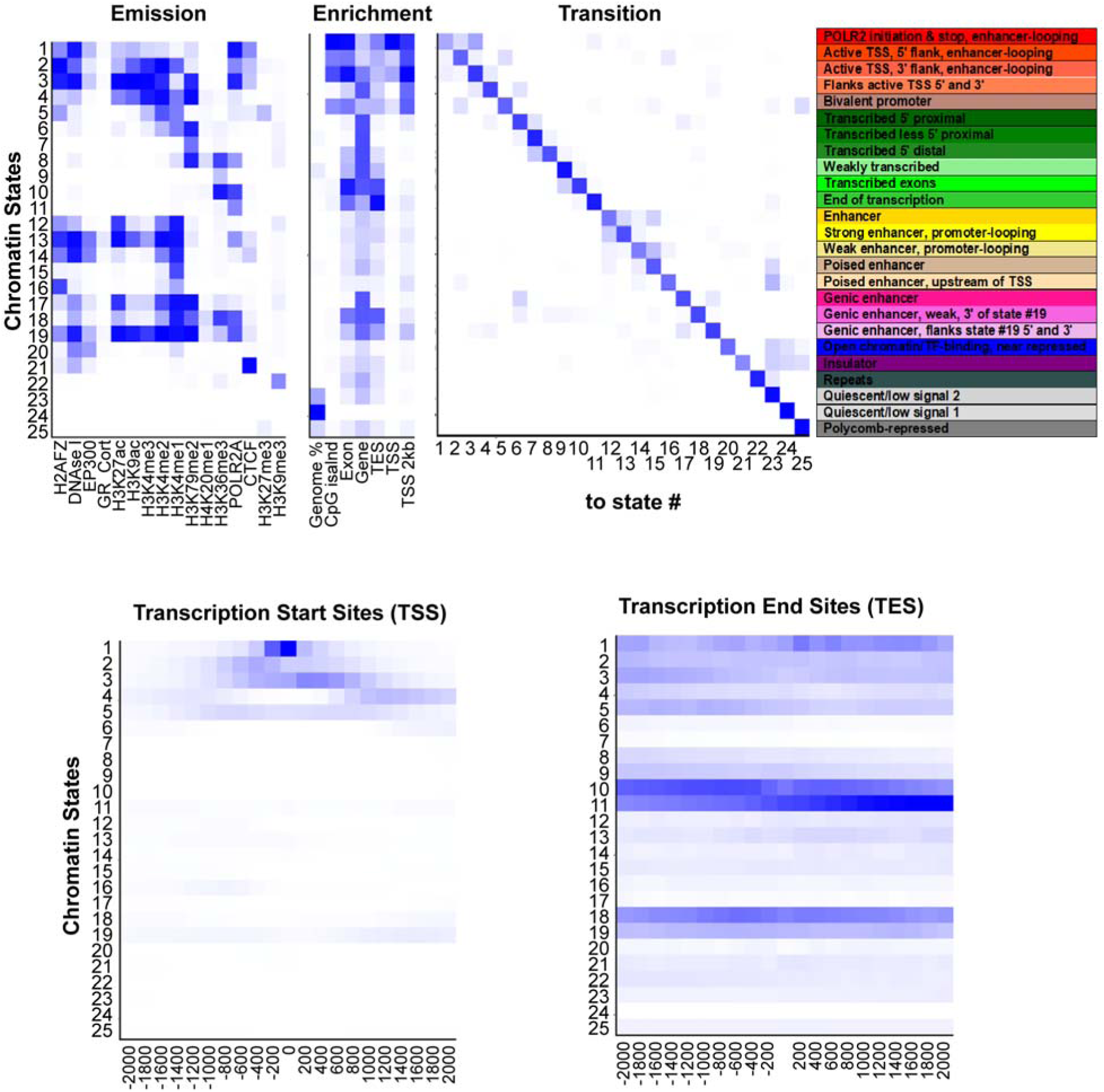
Output of ChromHMM analysis. The emission panel depicts the occurrence probability for each epigenetic mark in a particular state. The combinatorial occurrence of the marks is one of the pieces of information essential for chromatin state annotation. The enrichment panel represents enrichment for each state within a particular region in the genome, with a zoom-in focus on transcription start sites and transcription end sites as shown in the lower panels. The transition panel represents the spatial proximity of each state to each other.

**Extended Fig.3.**
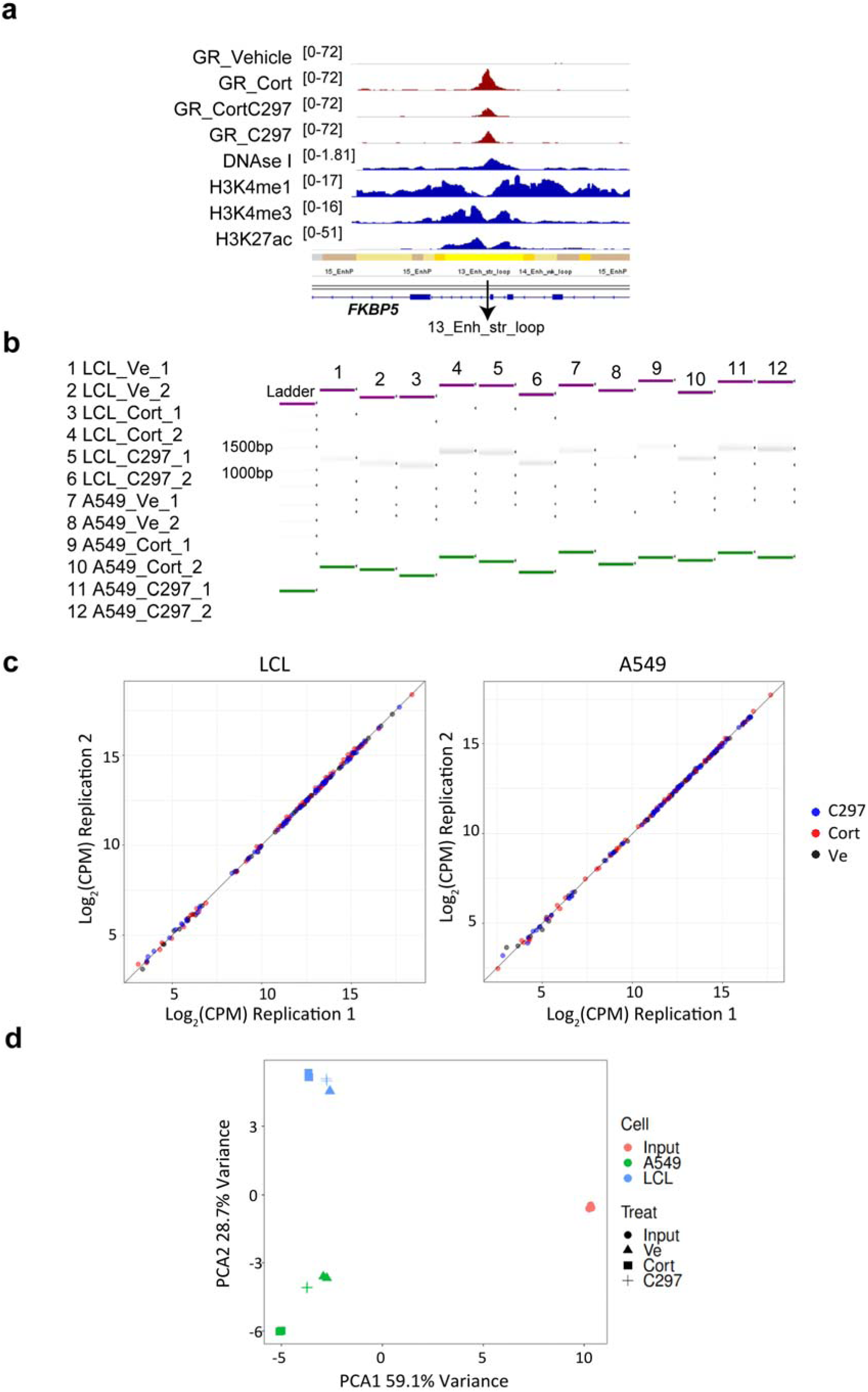
STARR-seq experimental design and statistical summary. (**a**) A glucocorticoid-induced GR peak that mapped to a strong enhancer region with looping properties at the 5’ end of *FKBP5* was included in the STARR-seq library as a positive control for enhancer activity. (**b**) After RNA transcribed from STARR-seq was enriched and converted to cDNA, a PCR reaction was conducted to transform cDNA to double-stranded DNA for sequencing library preparation. The image in (**b**) shows the STARR-seq DNA products measured by Agilent High Sensitivity D5000 ScreenTape, which appeared around 1300bp, the expected average size. (**c**) Correlation of STARR-seq read counts between 2 replicates showed good replication. (**d**) Principal component analysis of STARR-seq results showed global differences across treatments as well as between different cell lines. Cell types are color coded. Different shapes represent different treatments.

**Extended Fig.4.**
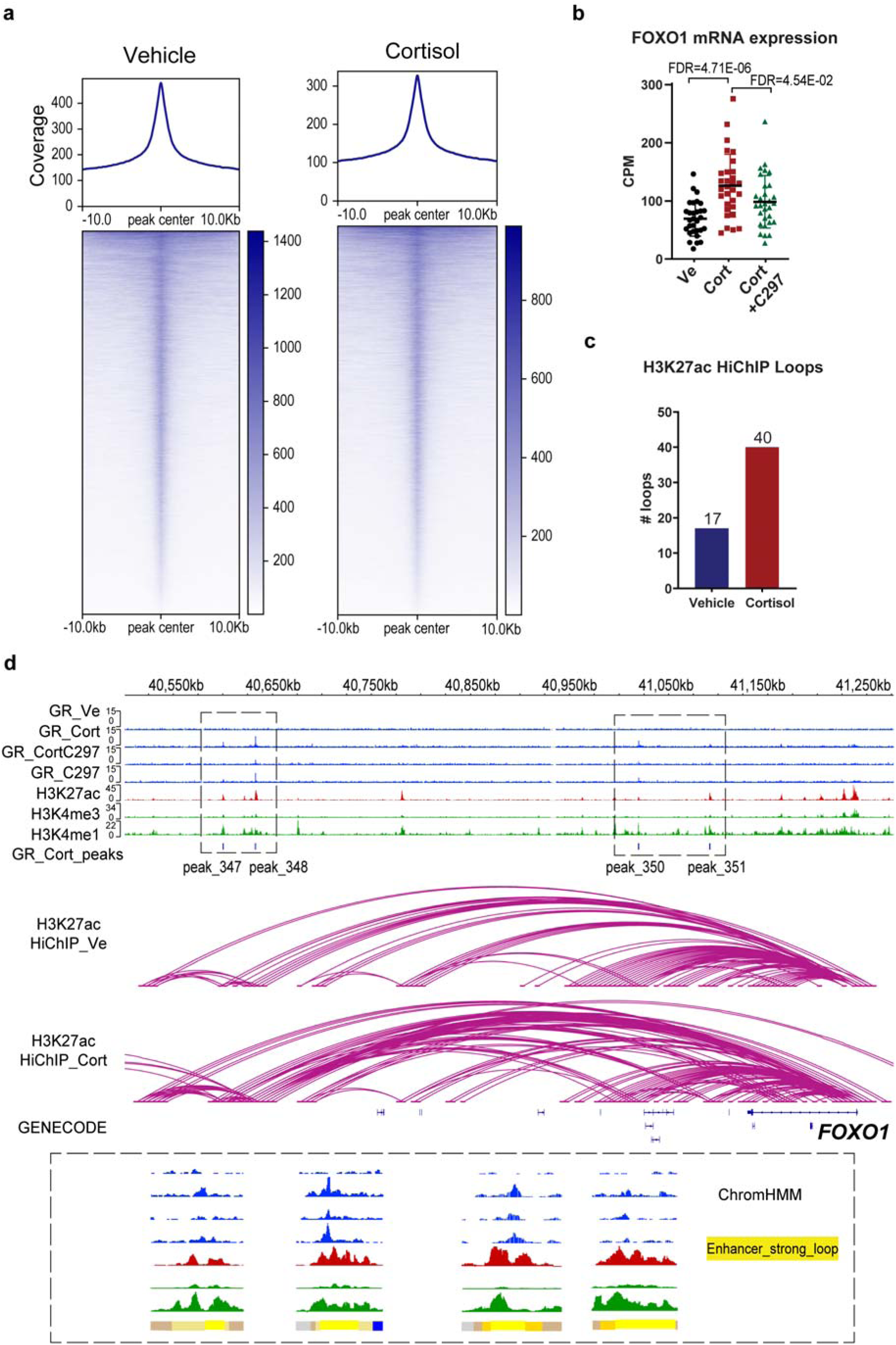
HiChIP experimental controls. (**a**) Enrichment of loops near known H3K27ac ChIP-peaks in the GM12878 LCL from ENCODE. (**b-d**) Additional positive control for “-omic” datasets after drug treatment showed their integration for another prototypical GR-targeted gene *FOXO1*. (**b**) mRNA expression of *FOXO1* before and after drug treatment as measured by RNA-seq. CPM stands for counts per million. (**c**) Number of HiChIP H3K27ac loops connecting GR-binding sites to *FOXO1* gene. (**d**) IGV plots of 4 different GR-dependent enhancers over a distance of 600kb, which together regulated *FOXO1*. These four enhancers were all predicted to be strong enhancers with looping properties by ChromHMM.

**Extended Fig.5.**
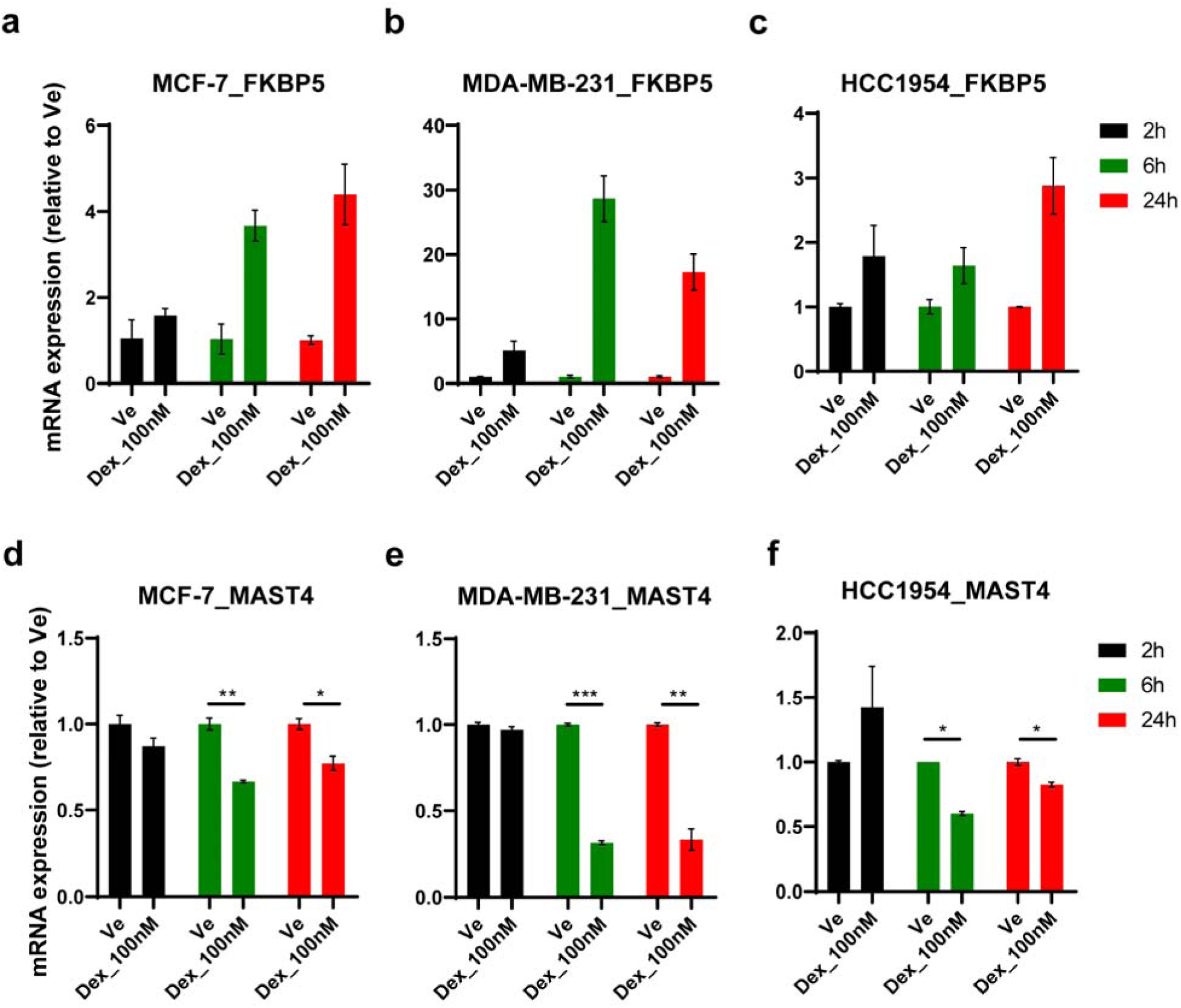
Glucocorticoid treatment of different breast cancer cell lines. The figures depict qPCR screening results for dexamethasone treatment of MCF-7 (ER+), MDA-MB-231 (TNBC), and HCC1954 (HER2+) breast cancer cell lines. *FKBP5* served as a positive control. * *P* < 0.05, ** *P* < 0.01, *** *P* < 0.001 from student’s t test.

**Extended Fig.6.**
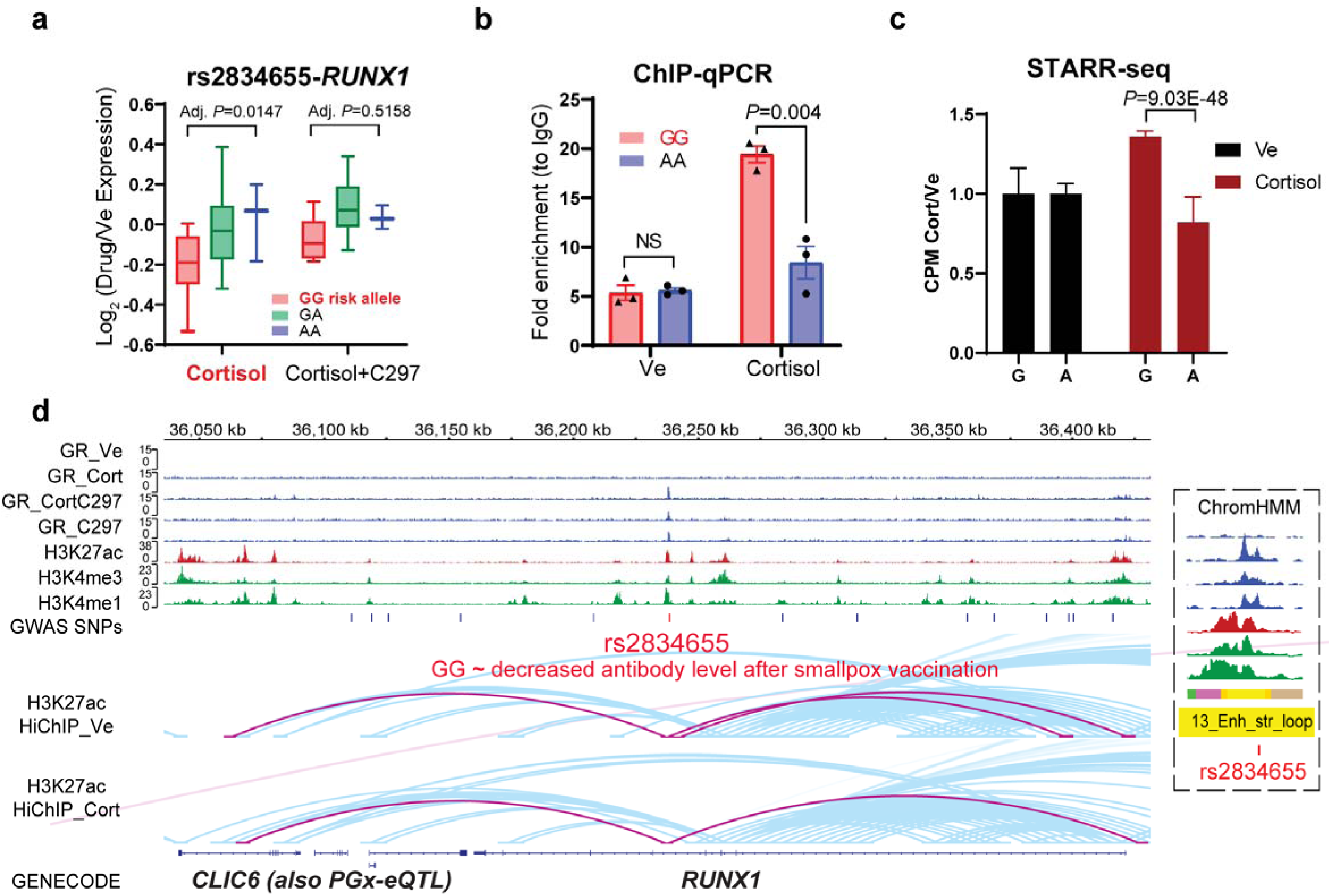
Example of a disease-associated PGx-eQTL SNP-gene pair for which the intronic enhancer SNP loops to the promoter of the eQTL gene in which it resides. (**a**) PGx-eQTL SNP-gene pair for rs2834655-*RUNX1*. Adjusted *P*-values from Tukey’s post-hoc multiple comparisons and represent differences between wildtype and variant genotypes. Cortisol induced the eQTL, and C297 antagonized the cortisol effect, normalizing eQTL expression. (**b**) ChIP-qPCR of GR demonstrating PGx SNP-mediated disruption of GR binding after cortisol treatment. This effect correlated with (**c**) lower enhancer activity as measured by STARR-seq. (**c**) SNP-dependent and drug-dependent enhancer activity of the PGx locus as measured by STARR-seq. CPM stands for counts per million. (**d**) IGV plots of the PGx SNP-eQTL gene locus. Tracks for GR-targeted ChIP-seq in different drug conditions are colored in blue, which show similar drug-dependent pattern as expression data: Cortisol induced GR binding at SNP locus, and C297 antagonized the cortisol effect, reducing GR binding. H3K4me1 is a histone mark associated with enhancers. H3K4me3 is a histone mark associated with promoters. H3K27ac is a histone mark associated with active promoters and enhancers. For the H3K27ac HiChIP tracks, loops directly interacting with the PGx SNP locus are highlighted in pink and others in blue.

**Extended Fig.7.**
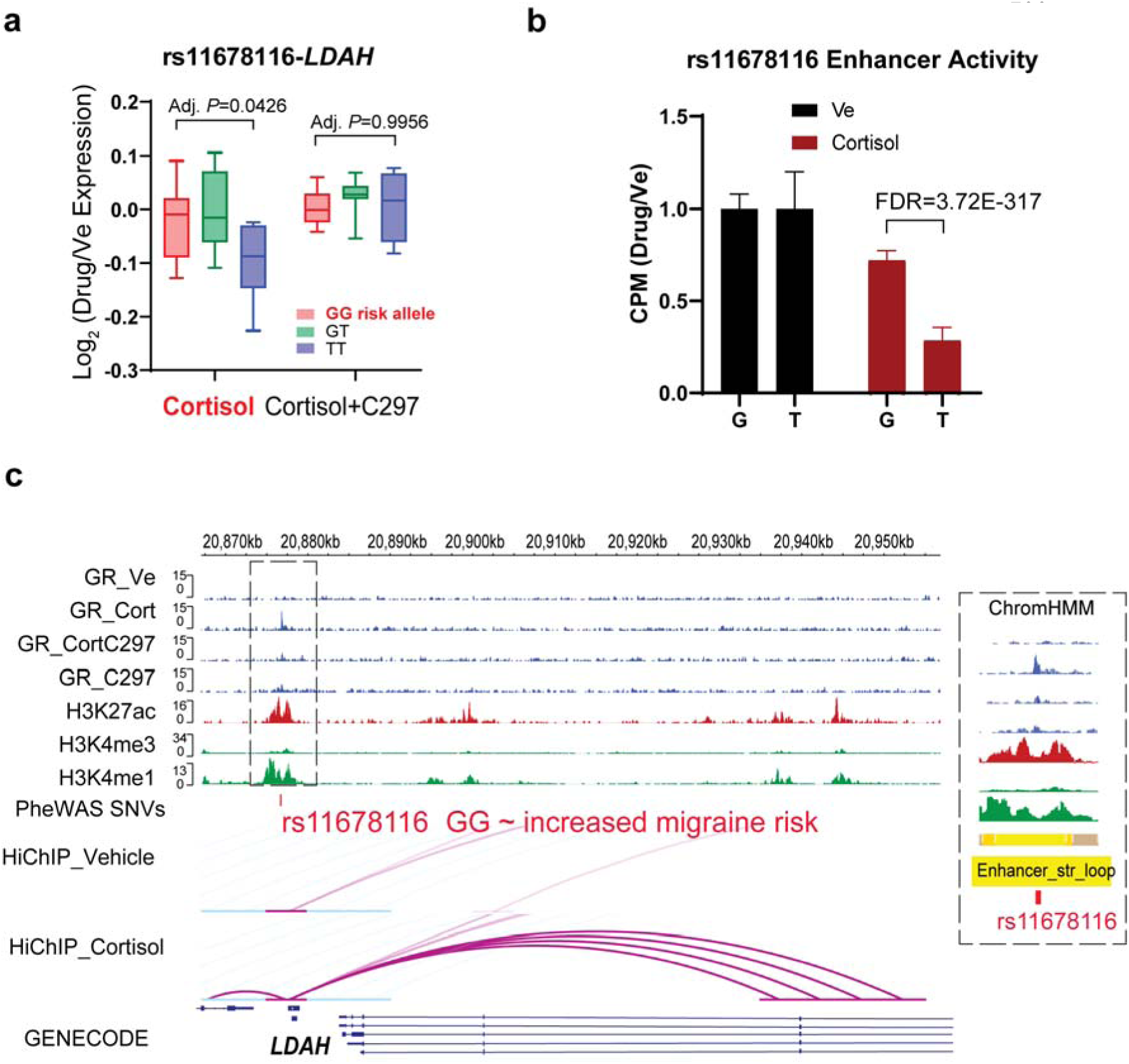
Example of a disease-associated PGx-eQTL SNP-gene pair for which induction of loops by cortisol was observed, bringing the intergenic enhancer SNP and the eQTL gene together. (**a**) PGx-eQTL SNP-gene pair for rs11678116-*LDAH*. Adjusted *P*-values from Tukey’s post-hoc multiple comparisons represent differences between wildtype and variant genotypes. Cortisol induced the eQTL, and C297 antagonized the cortisol effect, normalizing eQTL expression. (**b**) SNP-dependent and drug-dependent enhancer activity of the PGx locus as measured by STARR-seq. CPM stands for counts per million. (**c**) IGV plots of the PGx SNP-eQTL gene locus.

**Extended Fig.8.**
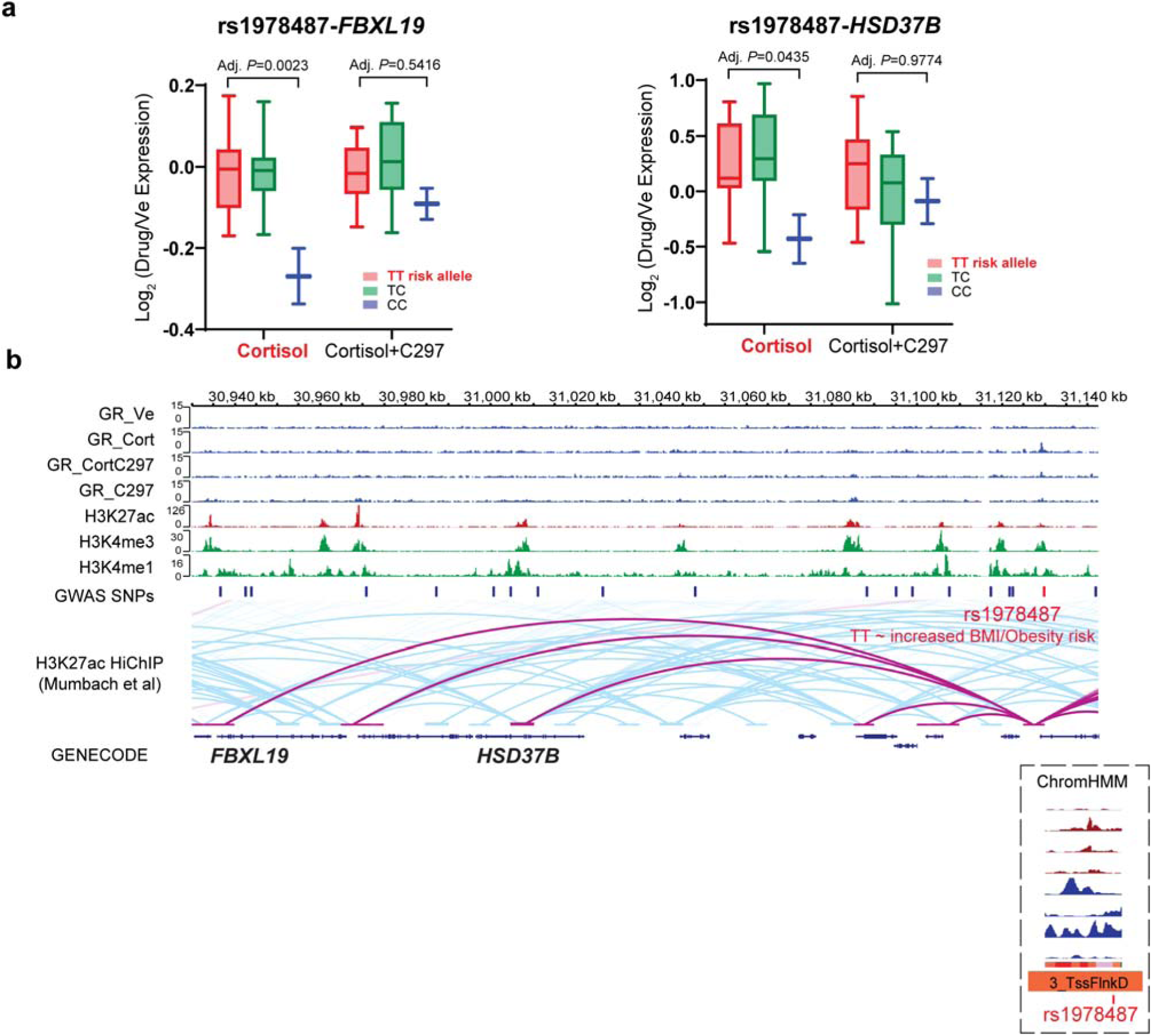
Example of a disease-associated PGx-eQTL SNP-gene pair for which the promoter SNP of a gene loops to the promoter of the eQTL genes located far away. (**a**) PGx-eQTL SNP-gene pairs for rs1978487-*FBXL19*/*HSD37B*. Adjusted *P*-values from Tukey’s multiple comparisons represent differences between wildtype and variant genotypes. Cortisol induced the eQTL, and C297 antagonized the cortisol effect, normalizing eQTL expression for both genes in similar fashion across genotypes. (**b**) IGV plots of the PGx SNP-eQTL gene locus.

## Methods

### Generation of SNP data

287 lymphoblastoid cell lines (LCLs) were obtained from the Coriell Institute. DNA from these 287 LCLs was genotyped with the Affymetrix Human SNP Array 6.0 at the Coriell Institute, and with the Illumina HumanHap550K and HumanExon510S-Duo Bead Chips in our laboratory. The genotype data were deposited in the National Center for Biotechnology Information Gene Expression Omnibus (GEO accession: GSE23120) ^54^.

### Cell culture & drug conditions

LCLs were cultured in RPMI 1640 supplemented with 15% FBS and 1% penicillin/streptomycin. A549 (ATCC^®^) cells were cultured in F12-K supplemented with 10% FBS and 1% penicillin/streptomycin. MCF-7 (ATCC^®^) cells were cultured in EMEM supplemented with 10% FBS and 1% penicillin/streptomycin. HCC1954 (ATCC^®^) cells were cultured in RMPI 1640 supplemented with 10% FBS and 1% penicillin/streptomycin. Cell culture conditions for these cell lines were 37°C and 5% CO2. MDA-MB-231 (ATCC^®^) cells were cultured in L-15 supplemented with 10% FBS. Cell culture condition for MDA-MB-231 were 37°C without CO2. Before glucocorticoid treatment experiments, all cells were grown in 5% charcoal-stripped media for 48 hours. Drug conditions and dosages were as follow: (1) Vehicle (dimethyl sulfoxide (DMSO) 0.1% and ethanol 0.1%), (2) hydrocortisone 100nM (Sigma Aldrich, dissolved in ethanol) plus DMSO 0.1%, (3) CORT108297 100nM (C297, AchemBlock, dissolved in DMSO) plus ethanol 0.1%, (4) hydrocortisone 100nM plus C297 100nM. Physiologically relevant dosages (less than 1000nM) and time points were optimized based on strongest induction of mRNA expression of GR-responsive canonical genes *FKBP5* and *GILZ* before saturation point.

### RNA-sequencing experiments

Thiry steroid-starved LCLs with similar expression of GR selected from the 300-LCL panel (**Table S1**) were subjected to the four drug exposure conditions in serum-free media for 9 hours. After treatment, cells were pelleted, and total RNA was extracted with the RNAeasy Mini Kit per manufacturer’s instruction (Qiagen). DNAse on-column treatment was performed with the DNase I set (Zymo). RNA sample integrity for all samples was 10. RNA-seq libraries were prepared with the TruSeq RNA Library Prep Kit v2 (Illumina). Paired-end sequencing 2x100bp was conducted on an Illumina HiSeq 4000 with a sequencing depth of ∼25 million paired-end reads per sample. Raw RNA sequencing reads were aligned to the human genome GRCh37 (hg19) using STAR ^55^. Raw counts were generated with the Python package “HTseq” ^56^ and normalized using conditional quantile normalization method (CQN) ^57^. Only genes that passed normalized counts of 32 in at least 15 cell lines and one drug condition were retained. Downstream differential expression analysis was conducted with the R package “EdgeR” ^58^ using a quasi-likelihood model.

### Glucocorticoid-targeted chromatin immunoprecipitation (ChIP)-sequencing experiment

Steroid-starved GM17261 cells were subjected to the four drug exposure conditions in serum-free media for 1 hour. 20 million cells for each condition were then cross-linked with 1% methanol-free formaldehyde (Thermo Fisher) for 10 minutes at room temperature. The reaction was then stopped with 125mM glycine for 5 minutes at room temperature and the pellets were frozen at -80°C prior to extraction. Chromatin preparation for ChIP-seq was performed as described by Zhong et al., 2017^59^. A cocktail of antibodies against the glucocorticoid receptor was added to chromatin input for immunoprecipitation: 2 μg of ab3570 (Abcam, lot GR3222141-5, discontinued) and 0.49 μg of 12041S (Cell Signaling Technology, lot 3) per 20 million cells. Please refer to Supplementary Methods for step-by-step details of the protocol. After library preparation, paired-end sequencing 2x50bp was performed on an Illumina HiSeq 4000 with sequencing depth of ∼12.5 million paired-end reads per sample. Raw sequencing reads were processed and analyzed using the HiChIP pipeline^60^ to obtain integrative genomics viewer files and a list of peaks (FDR < 0.01).

### Massively parallel reporter assay (STARR-seq) experiments

#### A. Input library construction

Human STARR-seq-ORI vector was obtained from Addgene (plasmid #99296). 8ug of STARR-seq plasmid was digested with 20uL AgeI-HF® (R3552, NEB), 20uL SalI-HF® (R3138, NEB), and 8uL Thermosensitive Alkaline Phosphatase (TSAP) in Multicore10X Buffer (Promega). Digestion was performed at 37 °C for 16 hours, followed by inactivation of TSAP at 74°C for 15 minutes. The digested products were then purified with QIAquick Gel Extraction Kit and Qiagen MinElute PCR purification kits. For inserts, we amplified all identified PGx-eQTL peaks extended by +/- 500bp using genomic DNA extracted from the 30 LCLs (**Table S1**). Primers were designed with the Nebuilder Assembly Tool v2.2.7. If the beginning or end of a region was difficult to anneal, we extended the region to include 50-100bp at the 3’ or 5’ end (for primers sequences and specific PCR conditions, refer to **Supplementary Data 4**). After amplification, every 10 PCR products were pooled together, gel purified, and cloned into the STARR-seq plasmid with NEBuilder® HiFi DNA Assembly Cloning Kit (E5520S, New England Biolabs) per manufacturer’s instruction. The ratio of vector (∼3000bp) to inserts (∼1200bp) was 0.05 pmol to 0.15 pmol. We included in the library a GR-induced peak within a strong enhancer region near the promoter of *FKBP5*, extended by +/-500bp, as a positive control. Assembled DNA products were then chemically transformed into NEB^®^ 5-alpha competent *E. coli* and spread on LB-ampicillin agar plates. The plates were scraped and grown in 2 liters of LB-ampicillin. Plasmids were extracted with the HiSpeed Plasmid Maxi Kit (Qiagen).

To check the quality of the inserted DNA loci (input library), 10ng plasmid DNA was amplified with KAPA HiFi HotStart ReadyMix PCR Kit (KK2601, KAPABiosystem) using as primers TTCTCTCCACAGGTGTCCA (forward) and GCAATAGCATCACAAATTTCACA (reverse) with the following conditions: 95°C for 3 minutes, followed by 9 cycles of 98°C for 20 seconds, 63°C for 15 seconds, and 72°C for 70 seconds. The PCR products were purified with QIAquick Gel Extraction Kit and Qiagen MinElute PCR purification kits and were then sequenced on a Hiseq4000 (Illumina) with 150-bp paired-end sequencing.

Due to their sensitivity to FBS, LCLs were not grown in charcoal-stripped media for this set of electroporation experiment as cell viability is a critical factor for successful STARR-seq screening. Therefore, to better evaluate GR activity on PGx loci cloned in STARR-seq plasmids, we also included A549, a lung cancer cell line in which GR has been extensively studied ^17^, in our STARR-seq screening. LCLs and A549 were maintained at viability of > 90% before transfection. For LCLs, 8 million cells were transfected with 20ug of plasmid DNA in one 100- uL cuvette using the Amaxa® Cell Line Nucleofector® Kit V (Lonza), program X-001. 10 independent pulses per drug condition were conducted, which resulted in a total of 600ug/240 million cells. For A549, 5 million cells were transfected with 10ug of plasmid DNA in one 100- uL cuvette using the Amaxa® Cell Line Nucleofector® Kit T (Lonza), program X-001. 15 independent pulses per drug condition were conducted, which resulted in a total of 450ug/225 million cells. After electroporation, cells were treated with drugs (hydrocortisone, C297 or vehicle) in 5% CS media for 9 hours. Total RNA was then harvested with the RNA Maxiprep kit per manufacturer’s instructions (Qiagen).

#### C. RNA library preparation and sequencing

Messenger RNA isolation was conducted with the Dynabeads Oligo (dT)_25_ (Invitrogen), followed by TURBO™ DNase digestion. The reaction was then cleaned up with RNA CleanXP beads (Beckman). mRNA concentrations were measured with Nanodrop 8000. We conducted first-strand cDNA synthesis with the SuperScript III Reverse Transcript kit (Invitrogen) using a reporter transcript-specific primer CTCATCAATGTATCTTATCATGTCTG and no more than 500ng mRNA per reaction. cDNA pool was then treated with RNAseA and purified with AMPure XP beads (Beckman). We conducted junction PCR with the KAPA HiFi HotStart ReadyMix PCR Kit (KK2601, KAPABiosystem) using primers TCGTGAGGCACTGGGCAG*G*T*G*T*C (forward) and CTTATCATGTCTGCTCGA*A*G*C (reverse) with the following conditions: 98°C for 45 seconds, followed by 16 cycles of 98°C for 15 seconds, 65°C for 30 seconds, and 72°C for 70 seconds. A total of 20 and 30 PCR reactions were conducted for LCLs and A549 cells, respectively. The PCR products were purified with AMPure XP beads (Beckman) before being sheared, adapter-ligated with NEBNext® Ultra™ DNA Library Prep Kit, and sequenced on an Illumina Hiseq 4000 with pair-end mode 2x150bp.

#### D. STARR-seq data analysis

Adapter sequences and reads aligned to universal plasmid sequences were trimmed out, and the trimmed reads were then aligned to GRCh37 (hg19) with BWA ^61^ mem using default parameters. Only reads with MAPQ ≥ 30 were kept for further analyses. Samtools ^62^ were used to convert sam files to bam files and to sort the bam files. Read counts for all SNPs sequenced within each PGx locus were then called with bcftools mpileup. Read counts of loci expression were called with the Python Package “HTseq” ^56^. Read counts for each sample were then normalized by library size and were log (CPM) transformed. Indels, multi-allelic variants, and variants with counts less than 20 were removed. All differential analyses were conducted with the R package “EdgeR” ^58^. For SNP-dependent loci analysis, the difference between the proportions of alternative alleles in vehicle- and drug-treated samples were evaluated using two-tailed Fisher’s exact test. Statistical significance was defined as FDR < 0.05, and % alternative allele difference (Alt_drug_/Total_drug_ – Alt_vehicle_/Total_vehicle_) > 10%. Visualization of STARR-seq data was conducted with the “EnhancedVolcano” R package (https://github.com/kevinblighe/EnhancedVolcano).

### Chromatin conformation capture of enhancer-enhancer/promoter interactions surrounding H3K27ac mark (HiChIP)

#### A. Publicly available data (baseline)

H3K27ac HiChIP data in GM12878 were downloaded from Mumbach et al., 2017 ^23^ and analyzed with the MAPS pipeline (48) using default parameters and known H3K27ac ChIP-seq peaks from ENCODE as anchors. All loops were called at 5kb bin size and were defined as having at least one end overlapping with a H3K27ac peak. PGx-eQTL SNPs were then overlapped with loop data to identify which SNP loci had H3K27ac-loop contact with eQTL genes using the R package “GenomicRanges”^63^.

#### B. Data generated in this study (before and after drug treatment)

Steroid-starved GM17261 cells was subjected to vehicle (0.01% EtOH) and 100nM of cortisol in serum-free media for 9 hours. After treatment, a portion of the cells was collected for RNA extraction and qRT-PCR to validate the effect of drug treatment on gene expression. 15 million cells per condition were fixed by 2% methanol-stabilized formaldehyde (Fisher Scientific, Cat #F79-500) at room temperature for 10 minutes. The reaction was then stopped with 125mM glycine for 5 minutes at room temperature. Cell pellets were snap-frozen in liquid nitrogen and sent to Arima Genomics (San Diego, CA, USA) for H3K27ac HiChIP library preparation. After passing quality control by shallow sequencing, the libraries were sequenced on an Illumina NovaSeq, yielding 500-800M pair-end reads per sample. HiChIP data were analyzed using the MAPS pipeline ^64^ using default parameters. Loops were called with a bin size of 5kb, maximum loop distance of 2000kb, and false discovery rate (FDR) of less than 0.01. Valid interacting pairs were defined as having at least one end overlapping with an H3K27ac peak. Two downstream analyses were conducted: (1) to identify loops before and after drug treatment which connected cortisol-dependent DEGs and cortisol ChIP-seq, (2) to identify loops before and after drug treatment which connected PGx SNPs and eQTL genes directly or through a common contact using the R package “GenomicRanges”^63^ and bedtools^65^. All genes were extended for 2000bp at the 5’ end (using ENSEMBL hg19 annotation) to include promoter regions.

### Pharmacogenomic-eQTL analysis

Analysis of eQTLs for normalized expression of drug/vehicle expression was conducted with the R package “Matrix eQTL”^66^ using ANOVA model. The number of SNP-gene pairs included in the eQTL analysis was narrowed down to leverage statistical power as described in the main text. Identified *P*-values of SNP-gene pairs after drug treatment were then compared with those obtained from analysis of 174 LCLs at baseline in the GTEx database with a significance cutoff of 0.05. PGx-eQTLs were also evaluated to determine whether the associations were lost after the antagonist treatment, demonstrating drug-dependent properties. SNPs in tight linkage disequilibrium (LD) (r^2^ > 0.4 and 0.8) with the identified PGx SNPs were investigated with regard to their potential to create/disrupt a known GR binding motif using the HaploReg v4.0 database. For significant SNP-gene pairs, we also conducted analyses of all SNPs within 200kb *cis* distance to compare the statistical significance of SNPs outside and inside GR binding sites, the results of which were plotted using the R package “Circlize”^67^. For selected examples depicted in the manuscript, post-hoc ANOVA Tukey’s test was used to evaluate significance for differences between homozygous wildtype and variant genotypes with GraphPad Prism 8.0. *P*-values were adjusted for multiple comparisons.

### Identification of pharmacogenomic-eQTL previously associated with a clinical phenotype

PGx-eQTL SNPs were overlapped with those documented in GWAS Catalog (https://www.ebi.ac.uk/gwas/), UKBiobank data (http://pheweb.sph.umich.edu/) or FinGen study data (https://r5.finngen.fi/about) to identify SNPs that have been associated with a clinical phenotype and whether the phenotype might fit with current knowledge about GR function. The *P-*values cutoff for GWAS and PheWAS were the same as those defined by the databases.

### ChromHMM analysis

Epigenomic datasets on GM12878 (LCL, Caucasian) were downloaded from the ENCODE project^20, 68^ for the following epigenetic marks/regulators: H3K4me1, H3K4me3, H3K27ac, H3K9me3, H3K27me3, H3K36me3, H3K4me2, H3K9ac, H4K20me1, H3K79me2, POLR2A, H2AFZ, DNAse hypersensitive sites, CTCF, and EP300^69^ (see **Supplemetary Data S3** for samples ID). BAM files of the LCL epigenetic marks/regulators, together with the BAM files of GR ChIP-seq after cortisol and C297 treatment, were binarized and segmented into 15, 18, or 25 chromatin states with default parameters using ChromHMM^19^. The 25-state value was adopted for its ability to capture as many states as possible without redundancy. These states were then annotated based on combinatorial and spatial patterns of chromatin marks^21^. Genomic coordinates of the chromatin states were converted from GRCh38/hg38 to GRCh19/hg19 using UCSC liftOver. Enrichment of PGx-eQTL SNPs in each of the 25 states was then analyzed using bedtools^65^.

### Functional validation of selected disease-associated PGx-eQTL experiments

#### Glucocorticoid-targeted ChIP-qPCR

To test GR activity for the rs1697139-*MAST4* PGx-eQTL locus in a breast cancer cell line, MDA-MB-231 cells were steroid-starved for 48 hours and subjected to vehicle and 100nM dexamethasone treatment for 1.5 hours (dexamethasone instead of cortisol was used to avoid cortisol effect on the mineralocorticoid receptor since, unlike LCLs, MDA-MB-231 expressed the mineralocorticoid receptor at reasonable levels). Similar ChIP-qPCR experiment conditions were conducted as described above. Two sets of primers targeting the rs1697139 GR-binding locus were used: Forward CACCTCAAAGGGTCCTCGGT, reverse TGTCACCACTTTCAGTTCCCTA (78 bp amplicon), and forward AGTGGTGACAACAGAGGATGTG, reverse TGTTCCTCGTCAGAAGCAGTT (79 bp amplicon).

To test SNP-dependent GR activity for rs12834655-*RUNX1* PGx-eQTL locus, GM17215 (rs12834655 genotype AA) and GM17293 (rs12834655 genotype GG) cells were steroid starved for 48 hours and subjected to vehicle and 100nM cortisol treatment for 1.5 hours. The chromatin immunoprecipitation steps and conditions were conducted as described above in the glucocorticoid-targeted ChIP-sequencing section. The Rabbit IgG control antibodies for ChIP-qPCR experiments were ab172730 (abcam, GR3298823-2, 2ug per 80ug chromatin), and Normal IgG Rabbit 2729 (Cell Signaling, lot 3, 0.49ug per 80ug chromatin). Quantitative PCR primers were designed with the NCBI Primers Blast Tools to ensure specificity of primers against the whole genome. The product size was 78bp, located inside the GR peak. The forward primer sequence was TTAGTCTTGACGTTGGCGGC, and reverse was GGTTCCAGCCGTGGCTTTAT. A primer targeting a GR-inducible peak upstream of *FKBP5* was used as a positive control similar to that in STARR-seq studies. The forward primer for this sequence was AGCGGGGGTTCTAGAGAGTG and reverse primer was CCAGGTTCTCAGGATCTCGT. The product size was 77bp. Each qPCR reaction was conducted with the *Power* SYBR™ Green PCR Master Mix (Thermo Fisher, Cat# 4367659) in triplicate. The Ct numbers of ChIP DNA were then normalized to Ct numbers of input DNA, and then normalized to Ct numbers of IgG. Data graphs were plotted using Prism (GraphPad Software). Statistical comparisons between genotypes were made using two-tailed student’s t-test.

### qRT-PCR of MAST4 in Breast Cancer Cell Lines after Glucocorticoids Treatment

To test the effect of GR signaling on *MAST4*, a gene that was a PGx-eQTL with the breast-cancer-associated rs1697139 SNP locus, 3 cell lines with reasonable expression of GR^70^ and MAST4 representing three common subtypes of breast cancer were subjected to treatment with 100nM Dexamethasone (Sigma, water soluble) for 2hrs, 6hrs, and 24hrs. Total RNA was extracted with Direct-zol RNA Miniprep (Zymo, Cat# R2052) per the manufacturer’s instructions. mRNA levels for *MAST4*, *GADPH*, and *FKBP5* were determined by qRT-PCR using the Power SYBR™ Green RNA-to-CT™ 1-Step Kit (Applied Biosystems Inc.). 100ng of total RNA was used for each reaction. Because MDA-MB-231 showed the most dramatic repression of *MAST4* after dexamethasone exposure, a dose-dependent 6-hour treatment with dexamethasone at 0nM, 1nM, 10nM, and 100nM was conducted on MDA-MB-231 cells, followed by qRT-PCR to confirm the drug effect. Analysis of qRT-PCR was conducted using the ^CT^ method^71^. Data graphs were then plotted using Prism (GraphPad Software). Statistical comparisons between genotypes were made using two-tailed student’s t-test.

### MAST4 Expression in Patient Tumor Samples

To assess *MAST4* expression in tumor samples from different subtypes of breast cancer as compared to normal tissue, data were downloaded from The Cancer Genome Atlas (TCGA). Two-tailed unpaired student’s t-test was used to compare *MAST4* expression between normal and cancer tissues. To assess *MAST4* expression and its relationship to relapse free survival (RFS) of breast cancer patients of all subtypes who underwent endocrine or chemotherapy, Kaplan-Meier plots were generated using KM plotter^72^ and its associated breast cancer database^73^. To address the concern that the association of *MAST4* expression with lower RFS was driven by its most significant repression in TNBC, which usually had a worse survival rate that other subtypes, Kaplan-Meier plots were also stratified by subtype, and *MAST4* low expression was predictive of lower RFS in other subtypes just as was the case with TNBC.

## References

1. MacArthur, J. et al. The new NHGRI-EBI Catalog of published genome-wide association studies (GWAS Catalog). Nucleic Acids Res 45, D896–D901 (2017).

2. Visscher, P.M. et al. 10 Years of GWAS Discovery: Biology, Function, and Translation. Am J Hum Genet 101, 5–22 (2017).

3. Consortium, G.T. The Genotype-Tissue Expression (GTEx) project. Nat Genet 45, 580–5 (2013).

4. Consortium, G.T. The GTEx Consortium atlas of genetic regulatory effects across human tissues. Science 369, 1318–1330 (2020).

5. Umans, B.D., Battle, A. & Gilad, Y. Where Are the Disease-Associated eQTLs? Trends Genet 37, 109–124 (2021).

6. Lee, M.N. et al. Common genetic variants modulate pathogen-sensing responses in human dendritic cells. Science 343, 1246980 (2014).

7. Strober, B.J. et al. Dynamic genetic regulation of gene expression during cellular differentiation. Science 364, 1287–1290 (2019).

8. Ingle, J.N. et al. Selective estrogen receptor modulators and pharmacogenomic variation in ZNF423 regulation of BRCA1 expression: individualized breast cancer prevention. Cancer Discov 3, 812–25 (2013).

9. Neavin, D.R. et al. Single Nucleotide Polymorphisms at a Distance from Aryl Hydrocarbon Receptor (AHR) Binding Sites Influence AHR Ligand-Dependent Gene Expression. Drug Metab Dispos 47, 983–994 (2019).

10. Liu, D. et al. TCF7L2 lncRNA: a link between bipolar disorder and body mass index through glucocorticoid signaling. Mol Psychiatry (2021).

11. Mangravite, L.M. et al. A statin-dependent QTL for GATM expression is associated with statin-induced myopathy. Nature 502, 377–80 (2013).

12. Arloth, J. et al. Genetic Differences in the Immediate Transcriptome Response to Stress Predict Risk-Related Brain Function and Psychiatric Disorders. Neuron 86, 1189–202 (2015).

13. Hunter, D.J. Gene-environment interactions in human diseases. Nat Rev Genet 6, 287–98 (2005).

14. Cain, D.W. & Cidlowski, J.A. Immune regulation by glucocorticoids. Nat Rev Immunol 17, 233–247 (2017).

15. Goodin, D.S. Glucocorticoid treatment of multiple sclerosis. Handb Clin Neurol 122, 455–64 (2014).

16. Mosca, M., Tani, C., Carli, L. & Bombardieri, S. Glucocorticoids in systemic lupus erythematosus. Clin Exp Rheumatol 29, S126–9 (2011).

17. Reddy, T.E. et al. Genomic determination of the glucocorticoid response reveals unexpected mechanisms of gene regulation. Genome Res 19, 2163–71 (2009).

18. Luca, F. et al. Genetic, functional and molecular features of glucocorticoid receptor binding. PLoS One 8, e61654 (2013).

19. Ernst, J. & Kellis, M. ChromHMM: automating chromatin-state discovery and characterization. Nature Methods 9, 215–216 (2012).

20. Consortium, E.P. An integrated encyclopedia of DNA elements in the human genome. Nature 489, 57–74 (2012).

21. Ernst, J. & Kellis, M. Large-scale imputation of epigenomic datasets for systematic annotation of diverse human tissues. Nat Biotechnol 33, 364–76 (2015).

22. Arnold, C.D. et al. Genome-Wide Quantitative Enhancer Activity Maps Identified by STARR-seq. Science 339, 1074–1077 (2013).

23. Mumbach, M.R. et al. Enhancer connectome in primary human cells identifies target genes of disease-associated DNA elements. Nat Genet 49, 1602–1612 (2017).

24. D’Ippolito, A.M. et al. Pre-established Chromatin Interactions Mediate the Genomic Response to Glucocorticoids. Cell Syst 7, 146–160 e7 (2018).

25. Michailidou, K. et al. Association analysis identifies 65 new breast cancer risk loci. Nature 551, 92–94 (2017).

26. Obradovic, M.M.S. et al. Glucocorticoids promote breast cancer metastasis. Nature 567, 540–544 (2019).

27. Pan, C. et al. Cisplatin-mediated activation of glucocorticoid receptor induces platinum resistance via MAST1. Nat Commun 12, 4960 (2021).

28. Sun, L. et al. [Identification of a novel human MAST4 gene, a new member of the microtubule associated serine-threonine kinase family]. Mol Biol (Mosk) 40, 808–15 (2006).

29. Neunert, C.E. Management of newly diagnosed immune thrombocytopenia: can we change outcomes? Blood Adv 1, 2295–2301 (2017).

30. Mevel, R., Draper, J.E., Lie, A.L.M., Kouskoff, V. & Lacaud, G. RUNX transcription factors: orchestrators of development. Development 146(2019).

31. Ovsyannikova, I.G. et al. Genome-wide association study of antibody response to smallpox vaccine. Vaccine 30, 4182–9 (2012).

32. Kobayashi, K.S. & van den Elsen, P.J. NLRC5: a key regulator of MHC class I-dependent immune responses. Nat Rev Immunol 12, 813–20 (2012).

33. Buckley, L. & Humphrey, M.B. Glucocorticoid-Induced Osteoporosis. N Engl J Med 379, 2547–2556 (2018).

34. Kennis, M. et al. Prospective biomarkers of major depressive disorder: a systematic review and meta-analysis. Mol Psychiatry 25, 321–338 (2020).

35. Belvederi Murri, M., et al. The HPA axis in bipolar disorder: Systematic review and meta-analysis. Psychoneuroendocrinology 63, 327–42 (2016).

36. Vegiopoulos, A. & Herzig, S. Glucocorticoids, metabolism and metabolic diseases. Mol Cell Endocrinol 275, 43–61 (2007).

37. Borsook, D., Maleki, N., Becerra, L. & McEwen, B. Understanding migraine through the lens of maladaptive stress responses: a model disease of allostatic load. Neuron 73, 219–34 (2012).

38. Rist, P.M., Tzourio, C. & Kurth, T. Associations between lipid levels and migraine: cross-sectional analysis in the epidemiology of vascular ageing study. Cephalalgia 31, 1459–65 (2011).

39. Acharya, A. et al. miR-26 suppresses adipocyte progenitor differentiation and fat production by targeting Fbxl19. Genes Dev 33, 1367–1380 (2019).

40. Rathjen, T. et al. Regulation of body weight and energy homeostasis by neuronal cell adhesion molecule 1. Nat Neurosci 20, 1096–1103 (2017).

41. Gamazon, E.R. et al. Using an atlas of gene regulation across 44 human tissues to inform complex disease- and trait-associated variation. Nat Genet 50, 956–967 (2018).

42. Gagliano Taliun, S.A., et al. Exploring and visualizing large-scale genetic associations by using PheWeb. Nat Genet 52, 550–552 (2020).

43. Vuckovic, D. et al. The Polygenic and Monogenic Basis of Blood Traits and Diseases. Cell 182, 1214–1231 e11 (2020).

44. Kichaev, G. et al. Leveraging Polygenic Functional Enrichment to Improve GWAS Power. Am J Hum Genet 104, 65–75 (2019).

45. Langefeld, C.D. et al. Transancestral mapping and genetic load in systemic lupus erythematosus. Nat Commun 8, 16021 (2017).

46. Andlauer, T.F. et al. Novel multiple sclerosis susceptibility loci implicated in epigenetic regulation. Sci Adv 2, e1501678 (2016).

47. Chen, M.H. et al. Trans-ethnic and Ancestry-Specific Blood-Cell Genetics in 746,667 Individuals from 5 Global Populations. Cell 182, 1198–1213 e14 (2020).

48. Matsunami, K. et al. Genome-Wide Association Study Identifies ZNF354C Variants Associated with Depression from Interferon-Based Therapy for Chronic Hepatitis C. PLoS One 11, e0164418 (2016).

49. Ward, J. et al. The genomic basis of mood instability: identification of 46 loci in 363,705 UK Biobank participants, genetic correlation with psychiatric disorders, and association with gene expression and function. Mol Psychiatry (2019).

50. Coleman, J.R.I. et al. The Genetics of the Mood Disorder Spectrum: Genome-wide Association Analyses of More Than 185,000 Cases and 439,000 Controls. Biol Psychiatry 88, 169–184 (2020).

51. Styrkarsdottir, U. et al. Multiple genetic loci for bone mineral density and fractures. N Engl J Med 358, 2355–65 (2008).

52. Wu, Y. et al. Genome-wide association study of medication-use and associated disease in the UK Biobank. Nature Communications 10(2019).

53. Vogelezang, S. et al. Novel loci for childhood body mass index and shared heritability with adult cardiometabolic traits. PLoS Genet 16, e1008718 (2020).

54. Niu, N. et al. Radiation pharmacogenomics: a genome-wide association approach to identify radiation response biomarkers using human lymphoblastoid cell lines. Genome Res 20, 1482–92 (2010).

55. Dobin, A. et al. STAR: ultrafast universal RNA-seq aligner. Bioinformatics 29, 15–21 (2013).

56. Anders, S., Pyl, P.T. & Huber, W. HTSeq--a Python framework to work with high-throughput sequencing data. Bioinformatics 31, 166–9 (2015).

57. Hansen, K.D., Irizarry, R.A. & Wu, Z. Removing technical variability in RNA-seq data using conditional quantile normalization. Biostatistics 13, 204–16 (2012).

58. Robinson, M.D., McCarthy, D.J. & Smyth, G.K. edgeR: a Bioconductor package for differential expression analysis of digital gene expression data. Bioinformatics 26, 139–40 (2010).

59. Zhong, J. et al. Purification of nanogram-range immunoprecipitated DNA in ChIP-seq application. BMC Genomics 18, 985 (2017).

60. Yan, H. et al. HiChIP: a high-throughput pipeline for integrative analysis of ChIP-Seq data. BMC Bioinformatics 15, 280 (2014).

61. Li, H. & Durbin, R. Fast and accurate short read alignment with Burrows-Wheeler transform. Bioinformatics 25, 1754–60 (2009).

62. Li, H. et al. The Sequence Alignment/Map format and SAMtools. Bioinformatics 25, 2078–9 (2009).

63. Lawrence, M. et al. Software for computing and annotating genomic ranges. PLoS Comput Biol 9, e1003118 (2013).

64. Juric, I. et al. MAPS: Model-based analysis of long-range chromatin interactions from PLAC-seq and HiChIP experiments. PLoS Comput Biol 15, e1006982 (2019).

65. Quinlan, A.R. & Hall, I.M. BEDTools: a flexible suite of utilities for comparing genomic features. Bioinformatics 26, 841–2 (2010).

66. Shabalin, A.A. Matrix eQTL: ultra fast eQTL analysis via large matrix operations. Bioinformatics 28, 1353–8 (2012).

67. Gu, Z., Gu, L., Eils, R., Schlesner, M. & Brors, B. circlize Implements and enhances circular visualization in R. Bioinformatics 30, 2811–2 (2014).

68. Davis, C.A. et al. The Encyclopedia of DNA elements (ENCODE): data portal update. Nucleic Acids Res 46, D794–D801 (2018).

69. Bernstein, B.E. et al. The NIH Roadmap Epigenomics Mapping Consortium. Nat Biotechnol 28, 1045–8 (2010).

70. Klijn, C. et al. A comprehensive transcriptional portrait of human cancer cell lines. Nat Biotechnol 33, 306–12 (2015).

71. Livak, K.J. & Schmittgen, T.D. Analysis of relative gene expression data using real-time quantitative PCR and the 2(-Delta Delta C(T)) Method. Methods 25, 402–8 (2001).

72. Gyorffy, B. Survival analysis across the entire transcriptome identifies biomarkers with the highest prognostic power in breast cancer. Comput Struct Biotechnol J 19, 4101–4109 (2021).

73. Gyorffy, B. et al. An online survival analysis tool to rapidly assess the effect of 22,277 genes on breast cancer prognosis using microarray data of 1,809 patients. Breast Cancer Res Treat 123, 725–31 (2010).

