## Supplementary Tables 1-3 for "Glucocorticoids Unmask Silent Non-Coding Genetic Risk Variants for Common Diseases"

Table S1. Information for the 30 LCLs used in this study

| **Cell ID** | **Catalog ID** | **Sex** | **Age (At sampling)** |
| --- | --- | --- | --- |
| CA01 | GM17201 | Male | 25 |
| CA07 | GM17207 | Male | 28 |
| CA12 | GM17212 | Male | 51 |
| CA14 | GM17214 | Male | 33 |
| CA15 | GM17215 | Female | 44 |
| CA18 | GM17218 | Female | 51 |
| CA21 | GM17221 | Female | 22 |
| CA36 | GM17236 | Female | 26 |
| CA38 | GM17238 | Female | 34 |
| CA40 | GM17240 | Male | 60 |
| CA41 | GM17241 | Male | 40 |
| CA45 | GM17245 | Female | 48 |
| CA49 | GM17249 | Male | 44 |
| CA50 | GM17250 | Female | 35 |
| CA56 | GM17256 | Female | 24 |
| CA58 | GM17258 | Female | 27 |
| CA60 | GM17260 | Female | 28 |
| CA61 | GM17261 | Female | 28 |
| CA63 | GM17263 | Female | 26 |
| CA64 | GM17264 | Male | 25 |
| CA65 | GM17265 | Female | 25 |
| CA66 | GM17266 | Female | 34 |
| CA68 | GM17268 | Male | 26 |
| CA69 | GM17269 | Male | 55 |
| CA79 | GM17279 | Female | 54 |
| CA84 | GM17284 | Female | 28 |
| CA86 | GM17286 | Female | 73 |
| CA89 | GM17289 | Female | 73 |
| CA93 | GM17293 | Female | 20 |
| CA94 | GM17294 | Female | 33 |

Table S2. Full annotation of 25 chromatin states built from 15 epigenetic marks in LCLs

| **STATE NO.** | **STATE ANNOTATION** | **ABBRIEVIATION** | **COLOR** | **COLOR CODE** |
| --- | --- | --- | --- | --- |
| 1 | POLR2 initiation & stop, enhancer-looping | TssTes_loop | Red | 255,0,0 |
| 2 | Active TSS, 5' flank, enhancer-looping | TssFlnkU_loop | Orange Red | 255,69,0 |
| 3 | Active TSS, 3' flank, enhancer-looping | TssFlnkD_loop | Tomato | 255,99,71 |
| 4 | Flanks active TSS 5' and 3' | TssFlnkUD | Coral | 255,127,80 |
| 5 | Bivalent promoter | TssBiv | Salmon | 250,128,114 |
| 6 | Transcribed 5' proximal | Tx1 | Darkgreen | 0,100,0 |
| 7 | Transcribed less 5' proximal | Tx2 | Green | 0,128,0 |
| 8 | Transcribed 5' distal | Tx3 | ForestGreen | 34,139,34 |
| 9 | Weakly transcribed | Tx4 | LightGreen | 144,238,144 |
| 10 | Transcribed exons | TxExon | Lime | 0,255,0 |
| 11 | End of transcription | TxEnd | Limegreen | 50,205,50 |
| 12 | Enhancer | Enh | Gold | 255,215,0 |
| 13 | Strong enhancer, promoter-looping | Enh_str_loop | Yellow | 255,255,0 |
| 14 | Weak enhancer, promoter-looping | Enh_wk_loop | Khaki | 240,230,140 |
| 15 | Poised enhancer | EnhP | Tan | 210,180,140 |
| 16 | Poised enhancer, upstream of TSS | EnhP_TssU | Navajowhite | 255,222,173 |
| 17 | Genic enhancer | EnhG1 | DeepPink | 255,20,147 |
| 18 | Genic enhancer, weak, 3' of state #17 | EnhG2 | LightPink | 255,182,193 |
| 19 | Genic enhancer, flanks state #17 5' and 3' | EnhG3 | Violet | 238,180,238 |
| 20 | Open chromatin/TF-binding, near repressed | OpenTF | Blue | 0,0,255 |
| 21 | Insulator | Ins | Purple | 128,0,128 |
| 22 | Repeats | Rpts | DarkSlateGray | 47,79,79 |
| 23 | Quiescent/low signal 1 | Quies | Lightgray | 211,211,211 |
| 24 | Quiescent/low signal 2 | Quies | Gainsboro | 220,220,220 |
| 25 | Polycomb-repressed | ReprPC | Gray | 128,128,128 |

Table S3. Distribution of PGx-eQTL SNPs in each chromatin state

**CORT**

| **Chromatin states** | **Counts of SNP-gene pair(s)** | |
| --- | --- | --- |
| 1_TssTes_loop | 1 | 1% |
| 10_TxExon | 1 | 1% |
| 12_Enh | 4 | 4% |
| 13_Enh_str_loop | 36 | 35% |
| 14_Enh_wk_loop | 6 | 6% |
| 15_EnhP | 11 | 11% |
| 16_EnhP_TssU | 1 | 1% |
| 17_EnhG1 | 6 | 6% |
| 18_EnhG2 | 6 | 6% |
| 19_EnhG3 | 12 | 12% |
| 20_OpenTF | 2 | 2% |
| 22_Rpts | 1 | 1% |
| 23_Quies | 7 | 7% |
| 25_ReprPC | 2 | 2% |
| 3_TssFlnkD_loop | 4 | 4% |
| 4_TssFlnkUD | 2 | 2% |
| Total | 102 | 100% |

**C297**

| **Chromatin states** | **Counts of SNP-gene pair(s)** | |
| --- | --- | --- |
| 13_Enh_str_loop | 19 | 59% |
| 14_Enh_wk_loop | 2 | 6% |
| 16_EnhP_TssU | 1 | 3% |
| 19_EnhG3 | 1 | 3% |
| 2_TssFlnkU_loop | 4 | 13% |
| 20_OpenTF | 1 | 3% |
| 3_TssFlnkD_loop | 4 | 13% |
| Grand Total | 32 | 100% |

Supplementary Data (Excel Sheet)

Data S1. (excel file) Differentially expressed genes after glucocorticoids treatment in 30 LCLs

Data S2. (excel file) Integrated information for all identified glucocorticoid-dependent PGx-eQTL SNP-gene pairs

Data S3. (excel file) ENCODE epigenetic datasets generated in LCL used for ChromHMM analysis

Data S4. (excel file) Primer sequences for STARR-seq locus library
